# Antibiotic acyldepsipeptides stimulate the *Streptomyces* Clp-ATPase/ClpP complex for accelerated proteolysis

**DOI:** 10.1101/2022.05.13.490424

**Authors:** Laura Reinhardt, Dhana Thomy, Markus Lakemeyer, Joaquin Ortega, Stephan A. Sieber, Peter Sass, Heike Brötz-Oesterhelt

**Affiliations:** Department of Microbial Bioactive Compounds, Interfaculty Institute of Microbiology and Infection Medicine, University of Tübingen, Auf der Morgenstelle 28, 72076 Tübingen, Germany; Cluster of Excellence - Controlling Microbes to Fight Infections, University of Tübingen, 72076 Tübingen, Germany; Department of Chemistry, Technical University of Munich, Lichtenbergstraße 4, 85748 Garching, Germany; Department of Anatomy and Cell Biology, McGill University, 3640 University Street, Montreal, Quebec H3A 0C7, Canada

**Keywords:** caseinolytic protease, AAA+ chaperones, Clp-ATPase, ClpP, ADEP, antimicrobial agents, mode of action

## Abstract

Clp proteases consist of a proteolytic, tetradecameric core ClpP and AAA+ Clp-ATPases. Streptomycetes, producers of a plethora of secondary metabolites, encode up to five different ClpP homologs and the composition of their unusually complex Clp protease machinery has remained unsolved. Here, we report on the composition of the house-keeping Clp protease in *Streptomyces*, consisting of a hetero-tetradecameric core built of ClpP1, ClpP2 and the cognate Clp-ATPases ClpX, ClpC1 or ClpC2, all interacting with ClpP2 only. ADEP antibiotics dysregulate the Clp protease for unregulated proteolysis. We observed that ADEP binds *Streptomyces* ClpP1, but not ClpP2, thereby not only triggering the degradation of non-native protein substrates but also accelerating Clp-ATPase-dependent proteolysis. The explanation is the concomitant binding of ADEP and Clp-ATPases to opposite sides of the ClpP1P2 barrel, hence revealing a third, so far unknown mechanism of ADEP action, i.e., the accelerated proteolysis of native protein substrates by the Clp protease.

**Significance:** Clp proteases are antibiotic and anti-cancer drug targets. Composed of the proteolytic core ClpP and a regulatory Clp-ATPase, the protease machinery is important for protein homeostasis and regulatory proteolysis. The acyldepsipeptide antibiotic ADEP targets ClpP and has shown promise for treating multi-resistant and persistent bacterial infections. The molecular mechanism of ADEP is multi-layered. Here, we present a new way how ADEP can deregulate the Clp protease system. Clp-ATPases and ADEP bind to opposite sides of *Streptomyces* ClpP, accelerating the degradation of natural Clp protease substrates. We also demonstrate the composition of the major *Streptomyces* Clp protease complex, a heteromeric ClpP1P2 core with the Clp-ATPases ClpX, ClpC1 or ClpC2 exclusively bound to ClpP2, and the killing mechanism of ADEP in *Streptomyces*.

## Introduction

Streptomycetes are known for their complex developmental life cycle as well as for the multitude of secondary metabolites they produce, including important marketed antibiotics. Morphological differentiation of the filamentous, multicellular bacteria is shaped by a variety of differentiation processes, which include the re-organization of chromosome segregation, cell division and cell-wall assembly, as well as the formation of a sporulating aerial mycelium (1). In addition, morphological differentiation is coordinated with the extraordinary diverse secondary metabolism. Such complex developmental programs often rely on the activation or inactivation of regulators that may be conferred by transcriptional control, protein modifications and/or regulated proteolysis. Energy-dependent degradation of short-lived regulators is one essential feature of the compartmentalized, tightly regulated bacterial caseinolytic protease Clp (2), which is intimately involved in the morphological differentiation of *Streptomyces* (3–6).

The Clp protease typically consists of a barrel-shaped, tetradecameric ClpP core that associates with hexameric ATP-consuming unfoldases, so-called Clp-ATPases, for protein degradation (7). The tetradecameric ClpP core is built of two stacked heptameric rings to form the proteolytic chamber of the Clp protease, secluding the catalytic site residues (Ser97, His122, Asp171 in *Escherichia coli*) within the inner lumen of the barrel, away from the cytoplasm (8). Access to the proteolytic chamber is tightly regulated, as substrate entry is only allowed via two axial pores, too small for protein passage, to prevent undesired substrate degradation. Consequently, in the absence of a cognate Clp-ATPase, ClpP lacks proteolytic activity and only degrades short peptides (9, 10). For regulated proteolysis, Clp-ATPases and associated adaptor proteins recognize natural Clp protease substrates, bind to the apical and/or distal surface of the ClpP core, and unfold and translocate the protein substrates under ATP consumption through the entrance pores into the proteolytic chamber of ClpP (11).

Many bacterial species encode only a single ClpP homolog, thus forming homo-tetradecameric ClpP complexes, for instance, the model bacteria *E. coli* and *Bacillus subtilis* or pathogens like *Staphylococcus aureus* and *Streptococcus pneumoniae* (8, 12–14). Other pathogenic bacteria, including mycobacteria, chlamydia, listeria, clostridia and pseudomonads, possess two ClpP isoforms, i.e. ClpP1 and ClpP2, which may form homo-tetradecameric complexes of either ClpP1 or ClpP2 or hetero-tetradecameric complexes involving both ClpP homologs (15–20). Streptomycetes have an even more complex Clp system and mostly encode three or five ClpP homologs. What is known about the *Streptomyces* Clp machinery is mainly based on whole-cell studies in *Streptomyces lividans*. In this species, five *clpP* genes are organized in two bicistronic (*clpP1clpP2* and *clpP3clpP4*) and one monocistronic (*clpP5*) operon (3, 21), whose expression was shown to be tightly controlled via several negative feedback loops (22–25). For example, the degradation of the transcriptional activators ClgR and PopR depends on the presence of ClpP1 and ClpP2 in *S. lividans* wild-type cells (Fig 1a). Here, ClgR induces the expression of ClpP1 and ClpP2, while the ClgR paralogue PopR is assumed to recruit the RNA polymerase for *clp3clpP4* transcription. As PopR was shown to be degraded by ClpP1 and ClpP2 in whole cells, PopR-mediated expression of the *clpP3clpP4* operon is silenced in the presence of these two ClpP homologs (21–24). On the other hand, ClpP3 and ClpP4 are expressed in *clpP1clpP2* deletion mutants due to the repealed degradation of PopR (21). In contrast to *E. coli* or *B. subtilis* where the function of ClpP is not essential for survival, at least one functional copy of either *clpP1clpP2* or *clpP3clpP4* is essential for viability of *S. lividans* (21). Thus, at least some functional redundancy seems to exist between the ClpP homologs in *Streptomyces* that might represent a rescue mechanism in case the function of ClpP1 and ClpP2 is disturbed or inhibited.

**Figure 1:**
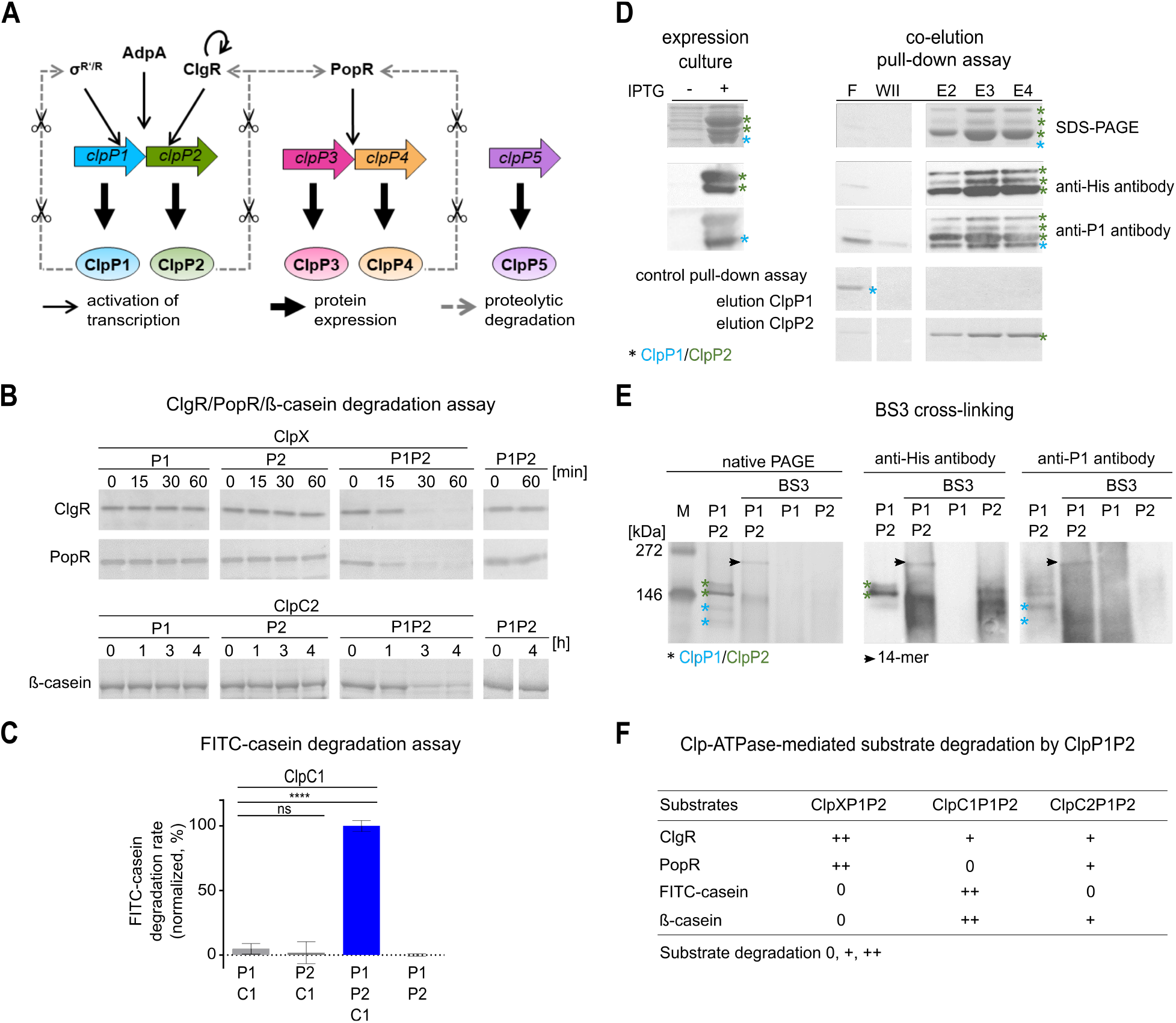
ClpP1 and ClpP2 form a hetero-tetradecameric core that interacts with the Clp-ATPases ClpX, ClpC1 and ClpC2 for protein degradation. **A.** Clp protease-dependent regulatory network in *Streptomyces*. **B** and **C.** *In vitro* protein degradation assays with purified Clp proteins indicate that the hydrolysis of protein substrates depends on the presence of ClpP1, ClpP2 and a partner Clp-ATPase. SDS-PAGE analyses of the degradation of the putative natural Clp substrates ClgR and PopR show that ClpX-mediated degradation of both substrates requires the presence of both ClpP1 and ClpP2 (B). Similarly, both ClpP1 and ClpP2 were required for ClpC2- or ClpC1-dependent digestion of ß-casein (B) or FITC-casein (C), respectively. The hydrolysis of FITC-casein was recorded as an increase in fluorescence intensity (relative fluorescence units, RFUs) over time. Mean values (normalized to %) of initial linear reaction kinetics are given. Statistical analyses were performed with one-way ANOVA by using three biological replicates, each comprising three technical replicates. P-values: ns > 0.05; **** ≤ 0.0001. Error bars indicate standard deviations. For SDS-PAGE images, representative examples of three replicates are shown. **D.** Co-elution of untagged ClpP1 and His_6_-tagged ClpP2 during Ni-NTA chromatography implies the interaction of the two ClpP homologs. Samples of the flow-through (F), the second wash fraction (WII) and the elution fractions E2-E4 are depicted. The cell lysate used for chromatography was generated after co-expression of ClpP1 (blue asterisk) and ClpP2-His_6_ (green asterisks) from two distinct, IPTG-inducible T7 promoters in the pETDUET-vector system. SDS-PAGE and immunoblot using specific anti-His_6_ or anti-ClpP1 antibodies. The three observed bands marked by the anti-His_6_ antibody correspond to full-length ClpP2 and processed variants (as discussed in a later section). Of note, the polyclonal anti-ClpP1 antiserum cross-reacted with ClpP2, which resulted in the detection of the same three bands as seen with the anti-His_6_ as well as an additional band that corresponds to ClpP1. Flow-through (F) and second wash fraction (W II) of the affinity chromatography experiment indicate the clearance of unbound ClpP1. In a control pull-down experiment, untagged ClpP1 and 6his-tagged ClpP2 were applied separately to a Ni-NTA column. Here, ClpP1 was only detected in the flow-through fraction (F), whereas ClpP2 was only detected in the elution fractions (E2-E4), thus proofing stringency of the pull-down assay. Assays were performed at least in triplicates, and representative SDS-PAGE images are shown. **E.** Cross-linking experiments visualize the heteromeric ClpP1P2 complex. Employing the chemical cross-linker BS3, a protein band with the approximate size of a ClpP tetradecamer appeared only in samples containing both ClpP1 and ClpP2. This band was detected with anti-ClpP1 as well as anti-His_6_ antibodies. A sample containing both ClpP1 and ClpP2 in the absence of BS3 did not yield the tetradecamer band, suggesting rather transient interactions between ClpP1 and ClpP2. Assays were performed at least in triplicates, and representative native PAGE images are shown. **F.** Overview of Clp-ATPase mediated substrate degradation experiments in combination with the proteolytic core ClpP1P2. Compare also degradation assays in Fig S1. 0, +, ++ reflect the efficiency of substrate degradation by the distinct Clp protease, respectively.

Despite the importance of the Clp system for streptomycetes, data on the exact molecular function of their Clp protease complex is scarce. In the current study, we set out to reconstitute and characterize the Clp system of *Streptomyces hawaiiensis* NRRL 15010 *in vitro*, which has the same *clpP* operon organization as *S. lividans* (26). *S. hawaiiensis* is of particular interest since it is the producer of antibiotic acyldepsipeptide 1 (ADEP1), the natural product progenitor of the potent class of ADEP antibiotics that bind to and deregulate the Clp protease complex (26–31).

We here present the molecular composition of the housekeeping Clp protease in *Streptomyces*, which consists of a hetero-tetradecameric ClpP1P2 core that interacts with corresponding Clp-ATPases solely via ClpP2. It is the first report on the full Clp protease machinery operating *in vitro*, including two ClpP isoforms and three Clp-ATPases working together to degrade natural Clp substrates as well as model substrates. In addition, our data assign a new, yet unprecedented mechanism of action to ADEP antibiotics, which is the accelerated degradation of natural Clp protease substrates as a result of the concomitant binding of ADEP and Clp-ATPase to opposite sides of the ClpP core.

## Results

### A hetero-tetradecameric ClpP1P2 complex interacts with ClpX, ClpC1 and ClpC2 for protein degradation

In order to investigate the composition and molecular function of the *Streptomyces* Clp protease *in vitro*, we first cloned, heterologously expressed and purified the *S. hawaiiensis* Clp proteins ClpP1 and ClpP2, their corresponding Clp-ATPase proteins ClpX, ClpC1 and ClpC2, as well as the predicted natural Clp protease substrates ClgR and PopR. We then performed *in vitro* degradation experiments by testing different combinations of ClpP and Clp-ATPase proteins to attempt the digestion of the natural substrates ClgR and PopR and the model protein substrates FITC-casein and ß-casein, respectively. Our data show that neither ClpP1 nor ClpP2 alone were capable of protein degradation in the presence of a Clp-ATPase (Fig 1BC). However, when ClpP1, ClpP2 and a Clp-ATPase were used in combination, protein substrates were rapidly digested, thus indicating that both ClpP1 and ClpP2 are required for Clp-ATPase-dependent protein degradation. Of note, not all protein substrates were degraded by all Clp-ATPase combinations or with similar efficiency. When we tested the degradation of ClgR and PopR by ClpP1 plus ClpP2 in the presence of either ClpX, ClpC1 or ClpC2, we observed that the presence of ClpX or ClpC2 leads to the degradation of both ClgR and PopR, although degradation was notably slower with ClpC2, whereas ClpC1 only conferred the degradation of ClgR, but not of PopR (Fig S1A). The model substrates ß-casein and FITC-casein were digested by ClpP1 plus ClpP2 in the presence of either ClpC1 or ClpC2, but not when ClpX was present (Fig S1BC).

Apparently, the stimulation of activity upon combining ClpP1 and ClpP2 points to the formation of a hetero-tetradecameric complex consisting of both ClpP homologs. To investigate a potential direct interaction of ClpP1 and ClpP2, we performed co-elution and native PAGE experiments using native ClpP1 and C-terminally His_6_-tagged ClpP2. During metal-ion affinity chromatography, ClpP1 was retained on the column by His_6_-tagged ClpP2 (Fig 1D), implying a direct interaction of ClpP1 and ClpP2. However, the interaction within the heteromeric ClpP1P2 complex seemed rather weak and the complex did not appear on standard native gels. To investigate this interaction further and substantiate the existence of a hetero-tetradecameric complex, we next performed cross-linking experiments using the chemical cross-linker BS3 to stabilize potential transient and weak interactions between both ClpP homologs. Now, the complex clearly showed and only the combination of both ClpP1 and ClpP2 allowed for the detection of a tetradecameric complex, as documented by native PAGE and immunoblotting analyses using anti-*Streptomyces* ClpP1 and anti-His_6_ antibodies (Fig 1E). In contrast, when ClpP1 or ClpP2 were used individually, no tetradecameric complexes were observed. Hence, our data show that the proteolytic core of the *Streptomyces* Clp protease consists of a ClpP1P2 hetero-tetradecameric complex that interacts with the Clp-ATPases ClpX, ClpC1 and ClpC2 for the degradation of protein substrates (Fig 1F).

### ClpP1 confers proteolytic activity to the complex, and ClpP2 interacts with the unfoldases ClpX, ClpC1 and ClpC2

Due to the observed heteromeric nature of the *Streptomyces* ClpP core, which consists of the two distinct homologs ClpP1 and ClpP2, we wondered whether both ClpP1 and ClpP2 equally contribute to Clp-ATPase-dependent proteolysis, or whether each homolog may fulfil certain, distinct functions during the degradation process. To approach this question, we first analysed the catalytic activity of the hetero-tetradecameric wild-type ClpP1P2 complex compared to combinations with the corresponding catalytic triad mutants ClpP1_S113A_ and ClpP2_S131A_, which had been constructed by replacing the nucleophilic serine moiety of the catalytic triad by an alanine residue via site-directed mutagenesis. Of note, both ClpP1 and ClpP2 of *S. hawaiiensis* comprise canonical Ser-His-Asp catalytic triads, similar to *S. lividans and S. coelicolor* (Fig S1D), but unlike ClpP1 from *Listeria monocytogenes. Listeria* ClpP1 features an uncommon asparagine residue instead of aspartate at position 172 that aligns to an active catalytic triad only when ClpP1 forms a complex with ClpP2 (32). Studying the turnover of the fluorogenic model substrate casein in the presence of *Streptomyces* ClpC1, we observed that a mixed core of ClpP1 and ClpP2_S131A_ retained full proteolytic activity at the level of wild-type ClpP1P2 (Fig 2A). This was in clear contrast to the mixed complexes ClpP1_S113A_P2 or ClpP1_S113A_P2_S131A_, in all of which protease activity was abolished, thus showing that the catalytic activity of the Clp complex is essentially conferred by ClpP1.

**Figure 2:**
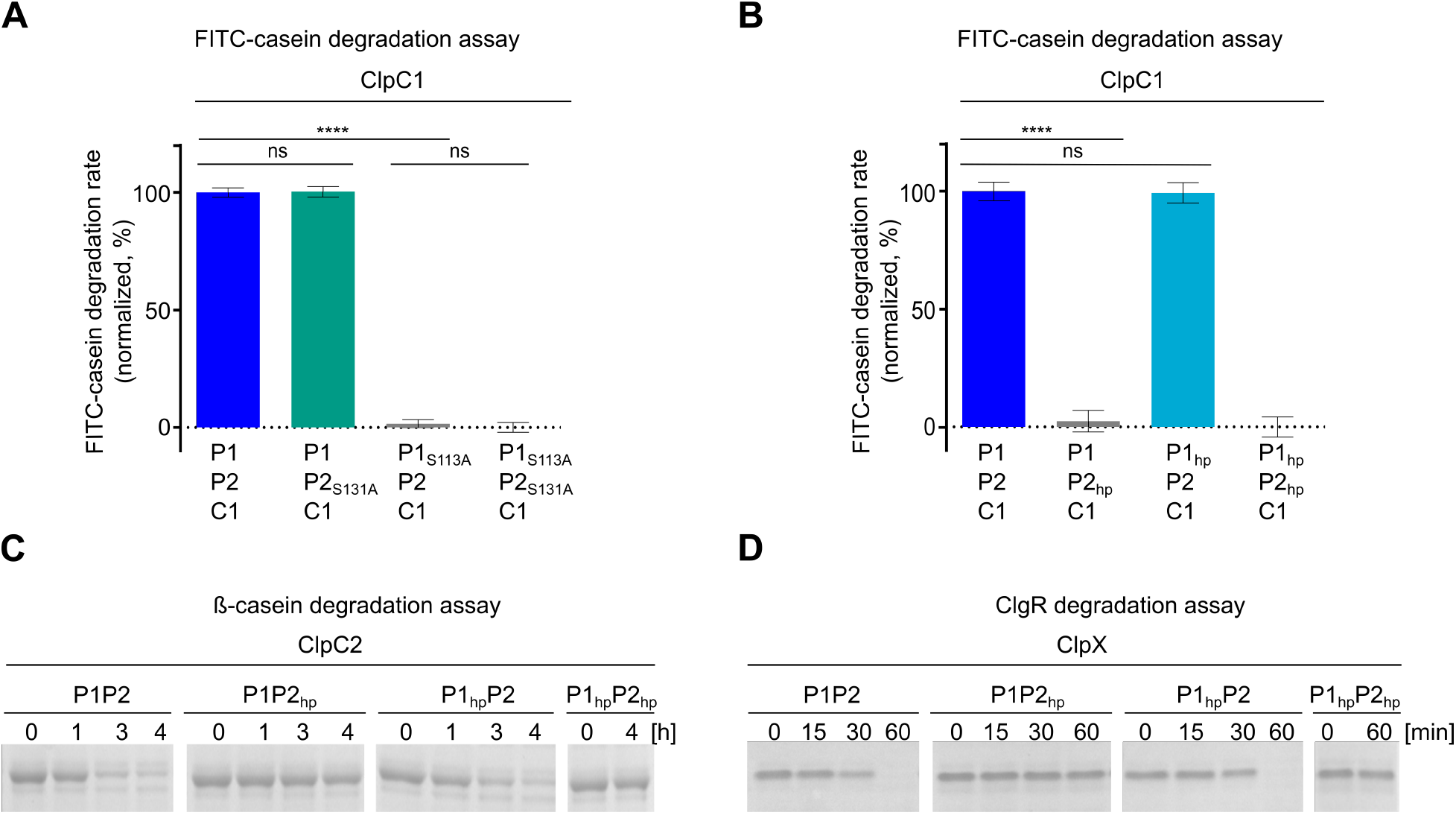
ClpP1 and ClpP2 have different but complementary functions during protein degradation. **A.** Impact of mutating the active site serine moieties in the ClpP1P2 hetero-complex. ClpP1_S113A_ but not ClpP2_S131A_ impairs ClpC1-mediated FITC-casein degradation, indicating that substrate protein hydrolysis is conferred by ClpP1. **B.** Effect of the hydrophobic pocket mutations in ClpP1_hp_ and ClpP2_hp_ on ClpC1-mediated FITC-casein degradation. Mutations in the hydrophobic pocket of ClpP2_hp_ prevent FITC-casein degradation, while for ClpP1_hp_ degradation is comparable to the wild-type protein, implying that the Clp-ATPase ClpC1 binds via ClpP2. For A and B, the hydrolysis of FITC-casein was recorded as RFU increase over time. Mean values (normalized to %) of initial linear reaction kinetics of the degradation curves are shown. P-values were calculated with one-way ANOVA using three biological replicates each comprising three technical replicates. P-values: ns > 0.05; **** ≤ 0.0001. Error bars indicate standard deviations. **C. and D.** Effects of the hydrophobic pocket mutations on ClpC2-mediated degradation of ß-casein (**C**) and ClpX-mediated degradation of ClgR (**D**). SDS-PAGE analyses show that protein degradation is inhibited in samples harbouring the hydrophobic pocket mutant ClpP2_hp_, whereas samples with ClpP1_hp_ retain wild-type activity, indicating that the Clp-ATPases ClpC2 and ClpX also interact via ClpP2. All assays were performed at least in triplicates and representative SDS-PAGE images are shown.

Aside from catalytic activity, the interaction with partner Clp-ATPases, which unfold and feed protein substrates into the ClpP degradation chamber, is a prerequisite for the proteolytic activity of the Clp protease. Therefore, we also probed the interaction of ClpP1 and ClpP2 with the cognate Clp-ATPases by generating particular hydrophobic pocket mutations, which had previously been shown to abrogate Clp-ATPase binding to the ClpP core in the related *M. tuberculosis* (MtClpP) (33). The corresponding three conserved aromatic residues (Y74, Y76, F96) in the crystal structure of *E. coli* ClpP (EcClpP) are also important for stabilizing Clp-ATPase interactions with ClpP (8), and the same positions were also shown to be involved in the binding of ADEP to *E. coli* and *B. subtilis* ClpP (34, 35). By aligning the protein sequences of *Streptomyces* ClpP1 and ClpP2 with EcClpP and MtClpP1P2 (Fig S1DE), we identified the corresponding amino acid positions in *Streptomyces* ClpP1 and ClpP2. To abrogate potential Clp-ATPase/ClpP interactions, but avoid substantial interference with protein structure and folding, suitable amino acids for exchange were selected according to the physicochemical properties and size of the amino acids, i.e., the bulky, aromatic tyrosine was replaced by a valine, whereas the smaller, hydrophilic serine was replaced by an alanine, yielding the hydrophobic pocket mutant proteins ClpP1_hp_ (Y76V, Y78V, Y98V) and ClpP2_hp_ (S94A, Y96V, Y116V) (Fig. S1DE). When using different combinations of wild-type and hydrophobic pocket mutant proteins of ClpP1 and ClpP2 in *in vitro* degradation assays, it emerged that protein substrates were only digested in the presence of wild-type ClpP2, no matter which of the three ATPases was tested (Fig 2B-D). Mutagenesis of the ClpP2 hydrophobic pocket residues resulted in complete inhibition of substrate turnover. In contrast, the corresponding mutations in ClpP1_hp_ did not affect protein degradation, and substrate turnover was comparable to the wild-type protein, indicating that the partner Clp-ATPases bind to the hetero-tetradecamer exclusively via ClpP2. Hence, our data clearly show that ClpP1 and ClpP2 of the hetero-tetradecameric ClpP core fulfil separate but complementary functions in the degradation process. ClpP1 confers catalytic activity to the complex, while ClpP2 is essential for the necessary interaction with the partner Clp-ATPases.

### ADEP antibiotics induce oligomerization and proteolytic activation of ClpP1, but not of ClpP2

*S. hawaiiensis* is the producer of the natural product antibiotic ADEP1 (Fig 3A) (26), and in a previous whole cell study with *S. lividans*, Mazodier and colleagues demonstrated that ClpP1 is targeted by this antibiotic (36). We therefore set out to further investigate the effect of ADEP antibiotics on the Clp system of the producer genus and to characterize the potential deregulation of the *in vitro* peptidase and protease activity of *S. hawaiiensis* ClpP1 and ClpP2 in the presence of ADEP1. To do so, we first used the fluorogenic dipeptide Suc-Leu-Tyr-aminomethylcoumarin (Suc-LY-AMC) as a substrate in peptidase activity assays and measured the fluorescence signal that results from peptide cleavage and concomitant AMC release. Our results showed that neither ClpP1, ClpP2 nor ClpP1P2 showed peptidase activity in the absence of ADEP1. However, ADEP1 clearly induced the hydrolysis of Suc-LY-AMC by either ClpP1 alone or by the mixed complex ClpP1P2, but not by ClpP2 (Fig 3BC). Corroborating our results above, peptidase activity was lost in all assays when ClpP1 was replaced by the active site mutant protein ClpP1_S113A_, while replacing ClpP2 by ClpP2_S131A_ had no effect on the activity of mixed complexes (Fig 3B). The same result was observed, when Z-GGL-AMC, Ac-WLA-AMC or Ac-Ala-hArg-2-Aoc-ACC were used as substrates in peptidase activity assays (Fig 3D). Probing the binding site, ADEP1 failed to induce peptide hydrolysis when ClpP1 was exchanged by the hydrophobic pocket mutant protein ClpP1_hp_, whereas replacement of ClpP2 by ClpP2_hp_ had no effect on substrate hydrolysis by the mixed complex (Fig 3B). These results indicate that ADEP1 interacts with ClpP1, but not ClpP2. Such observed asymmetric binding was not restricted to ADEP1 but was also detected for ADEP2 and ADEP4, two synthetic derivatives previously optimized for potent anti-*S. aureus* activity (27). Uniformly, all ADEPs induced peptidase activity via binding to ClpP1, but not to ClpP2 (Fig 3BE). Interestingly, ClpP1P2 exhibited slightly higher peptidase activity when activated by ADEP1, as compared to ADEP2 and ADEP4, thereby indicating that the binding of the natural compound ADEP1 to *Streptomyces* ClpP1 may have been evolutionary optimized. Mixed ClpP1P2, ClpP1P2_S131A_ and ClpP1P2_hp_ exhibited superior peptidase activity in the presence of ADEP1 compared to ClpP1 alone (Fig 3B), implying a stimulation of ClpP1 catalytic activity by the interaction with ClpP2. Of note, the presence of ADEP-ClpP1P2 complexes was verified by native PAGE and size exclusion chromatography (Fig S2A-E). Similar results were observed in protease activity assays using fluorogenic FITC-casein as a substrate and measuring FITC release upon casein degradation (Fig 3F). Also, here, ADEP1 induced proteolysis of casein by either ClpP1 alone or by mixed ClpP1P2, but not by ClpP2 and again the mixed complex performed better than ClpP1 alone. Thus, our data reveal that ADEP1 activates the purified and otherwise dormant *Streptomyces* ClpP core to Clp-ATPase-independent peptidase and protease activity by binding to and deregulating the activity of ClpP1.

**Figure 3:**
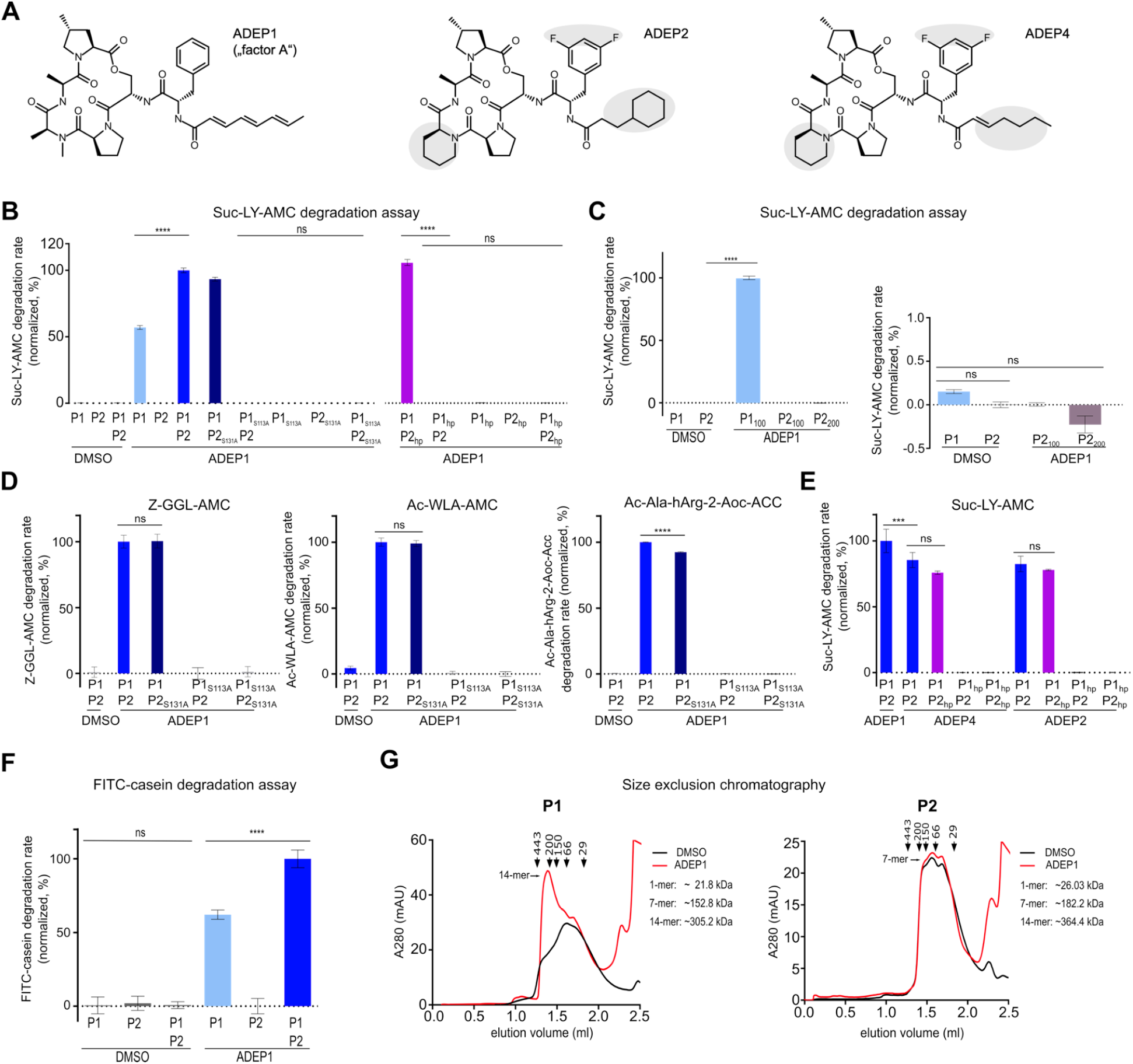
ADEP binds to and deregulates ClpP1, but not ClpP2. **A.** Chemical structures of the natural product ADEP1 and its synthetic congeners ADEP2 and ADEP4. ADEP1, representing “factor A” of the A54556 antibiotic complex produced by *S. hawaiiensis* NRRL 15010, consists of a macrolactone core, an aliphatic side chain, and an *N*-acylphenylalanine linker. Differences in the chemical structures between these ADEP derivatives are indicated in grey. **B-E**. Effect of ADEP on the peptidase activity of ClpP1 and ClpP2. Peptidase activity assays using the fluorogenic dipeptide Suc-LY-AMC as substrate show that in the absence of ADEP1 (DMSO controls), ClpP1 and ClpP2 alone or combined are devoid of peptidase activity. **B.** Addition of ADEP1 triggered peptidase activity of ClpP1 alone and even stronger of the ClpP1P2 hetero-complex, while ClpP2 alone remained inactive. Peptide hydrolysis was prevented in samples containing the catalytic triad or hydrophobic pocket mutant proteins ClpP1_S113A_ or ClpP1_hp_, respectively, but not in the corresponding ClpP2 mutants, thus indicating ADEP1 binding via ClpP1. The slopes of initial degradation rates are shown, normalized in %. **C.** Elevated concentrations of ADEP1 (100 μM or 200 μM) did not lead to detectable peptidase activity of ClpP2 (4 μM). ClpP1 served as a positive control. The slopes of initial degradation rates are shown (normalized in %; ClpP1 activity set to 100%). The right panel is the magnified area near the base line (Y-axis from −0.5 to 1%). ClpP1 activity without ADEP is only 0.15% of the ClpP1 activity in presence of ADEP. **D.** Peptidase assays using the fluorogenic peptide substrates Z-GGL-AMC, Ac-WLA-AMC and Ac-Ala-hArg-2-Aoc-Acc confirm that peptidase activity is mediated by ClpP1, but not by ClpP2. **E.** Peptidase assays using the derivatives ADEP2 and ADEP4 in comparison to ADEP1. The data indicates that asymmetric binding to ClpP1 is not restricted to ADEP1 but occurs also for ADEP2 and ADEP4. Of note, ClpP1P2 exhibited slightly higher peptidase activity in the presence of ADEP1 compared to ADEP2 and ADEP4. **F.** Effect of ADEP1 on the protease activity of ClpP1 and ClpP2 using FITC-casein as a substrate. ADEP1 activated casein degradation by ClpP1 alone and even stronger by the ClpP1P2 hetero-complex, while ClpP2 alone remained inactive. DMSO was used in control reactions. Mean values (normalized to %) of initial linear reaction kinetics are given. In B-F, error bars represent standard deviations. Statistical analyses were performed with one-way ANOVA using three biological replicates each comprising three technical replicates, except for the Suc-LY-AMC degradation assays with the hydrophobic pocket mutants ClpP1_hp_ and ClpP2_hp_, where two biological replicates were used. P-values: ns > 0.05; ****≤ 0.0001. Error bars indicate standard deviations. **G.** Oligomeric state analyses of ClpP1 and ClpP2 as determined by size exclusion chromatography. ADEP1 triggered the assembly of ClpP1 into homo-tetradecamers, which explains the independent peptidase and protease activity of ADEP1-activated ClpP1. In contrast, ClpP2 did not form homo-tetradecamers. DMSO was used as a control. Depicted experiments are representatives of three biological replicates.

Next, we analysed the oligomeric states of ClpP1 and ClpP2 in the absence and presence of ADEP1 by size exclusion chromatography (Fig 3G). Regarding ClpP1, it emerged that the addition of ADEP1 clearly led to a distinct shift from lower oligomeric states to ClpP1 tetradecamers (calculated molecular mass of 305.2 kDa). In contrast, the elution profile of ClpP2 did not change in the presence of ADEP1 and ClpP2 mainly eluted as heptamers (calculated molecular mass of 182.2 kDa) or lower oligomeric species, while ClpP2 tetradecamers (calculated molecular mass of 364.4 kDa) were not detected (Fig 3G). Hence, our data show that ADEP1 binds to ClpP1 and triggers its oligomerization into catalytically active ClpP1 tetradecamers. This explains the independent peptidase and protease activity of ClpP1 in the presence of ADEP1, while ClpP2 does not appear to interact with ADEP1 under the conditions tested.

In line with this, it is noteworthy that ClpP1 and ClpP2 proteins underwent processing reactions in the course of our *in vitro* degradation experiments (Fig S3, S4). *In vitro*, processing of ClpP1 relied either on the binding of ClpXP2 or on the binding of ADEP (regardless of ClpP2), while processing of ClpP2 strictly depended on the presence of ClpP1, but not on ClpX or ADEP (Fig S3AB). However, processing of both ClpP1 and ClpP2 fully relied on the integrity of the catalytic triad of ClpP1 (Fig S3C) and did not alter the requirement of hetero-tetradecameric complexes for proteolytic activity (Fig S4F), thereby corroborating our hitherto obtained results.

### Differences in the binding pocket of ClpP1 versus ClpP2 modulate specificity to ADEP and ClpC1

Since ADEP and Clp-ATPases usually bind to the same hydrophobic pockets of the ClpP barrel, it is intriguing that ClpP1 emerged ADEP-sensitive but did not interact with Clp-ATPases, whereas ClpP2 was ADEP-insensitive while constituting the main interaction partner for ClpX, ClpC1 and ClpC2. When analysing the hydrophobic pockets of ClpP1 and ClpP2, it became apparent that both isoforms differ in the first amino acid residue that was selected for mutation of each ClpP hydrophobic pocket (Y76 in ClpP1, S94 in ClpP2, see above). Since tyrosine and serine obviously differ regarding their size and physicochemical properties, we hypothesized that this difference may influence the binding capability of ClpP1 versus ClpP2 for ADEP and corresponding Clp-ATPases. We therefore exchanged Y76 of ClpP1 by a serine and, vice versa, S94 of ClpP2 by a tyrosine, thereby yielding the hydrophobic pocket mutant proteins ClpP1_Y76S_ and ClpP2_S94Y_. Native PAGE showed that ClpP1_Y76S_ was notably impaired in ADEP-induced assembly of homo-tetradecamers (Fig S2F), and consequently, in the presence of ADEP, ClpP1_Y76S_-containing complexes showed substantially decreased peptidase activity compared to wtClpP1 (Fig S2G), together indicating that the binding of ADEP to ClpP1_Y76S_ was impaired. Of note, ClpC1-mediated protease activity of ClpPl_Y76S_P2 was also reduced, suggesting that Y76 may also have a role in Clp protease assembly and/or catalysis. Now, probing ClpP2_S94Y_, protease activity of ClpC1P1P2_S94Y_ was significantly reduced, suggesting impaired interaction of ClpC1 with ClpP2_S94Y_. However, the ClpP2_S94Y_ mutation did not suffice for making ClpP2 responsive to ADEP (peptidase activity could not be activated by ADEP). Thus, our results clearly indicate that Y76 in ClpP1 and S94 in ClpP2 play crucial roles in conferring the specificity to ADEP versus ClpC1, respectively. However, switching Y76 in ClpP1 and S94 in ClpP2 was not sufficient to also switch the individual binding specificities regarding ADEP and ClpC1 entirely, which may be due to the individual protein characteristics of both ClpP1 and ClpP2, e.g., intramolecular bonds, folding characteristics, cavity spacing, and/or further potentially contributing amino acids.

### Whole cell studies corroborate the existence of a heteromeric, asymmetric Clp protease complex in *Streptomyces*

Our study so far employed purified *S. hawaiiensis* proteins *in vitro* to elucidate the molecular composition and function of the *Streptomyces* Clp protease. To verify our findings on the whole cell level and to also transfer these findings to other *Streptomyces* strains, we chose *S. lividans* as a model strain for confirmative whole cell studies, since earlier studies on the Clp system in streptomycetes were focussed on this organism (21–23, 26, 36). In this context, it is noteworthy, that ClpP1 and ClpP2 of *S. hawaiiensis, S. lividans* and *S. coelicolor* appear closely related and share high sequence similarity (Fig S1D). To this end, we constructed in-frame Δ*clpP1* (Δ*SlP1*) as well as Δ*clpP1clpP2* (Δ*SlP1P2*) deletion mutants in *S. lividans*, which we complemented with either *SlclpP1* or *SlclpP1P2*, respectively. In addition, C-terminally His_6_-tagged SlClpP2 (SlClpP2-His_6_) was constructed for detection purposes. Cell extracts of the respective deletion and complemented mutants were then analysed via immunoblotting for the SlPopR-dependent expression of SlClpP3 by utilizing anti-ClpP1, anti-ClpP3, and anti-His_6_ antibodies. Here, we interpreted the absence of SlClpP3 as an indicator for the functional state of the SlClpP1P2 system, capitalizing on previous studies in *S. lividans* (22), which had shown that the degradation of SlPopR by SlClpP1P2-mediated regulatory proteolysis prevents the expression of SlClpP3. In short, whenever SlClpP3 is present, SlClpP1P2 is not capable of degrading SlPopR (Fig 4A). In line with our *in vitro* results described above, *in vivo* SlClpP1P2 protease activity (and corresponding absence of SlClpP3) could only be observed, when both SlClpP1 and SlClpP2 were present in the cell (Fig 4B, S5A), thus underlining the requirement of a functional interaction of ClpP1 and ClpP2.

**Figure 4:**
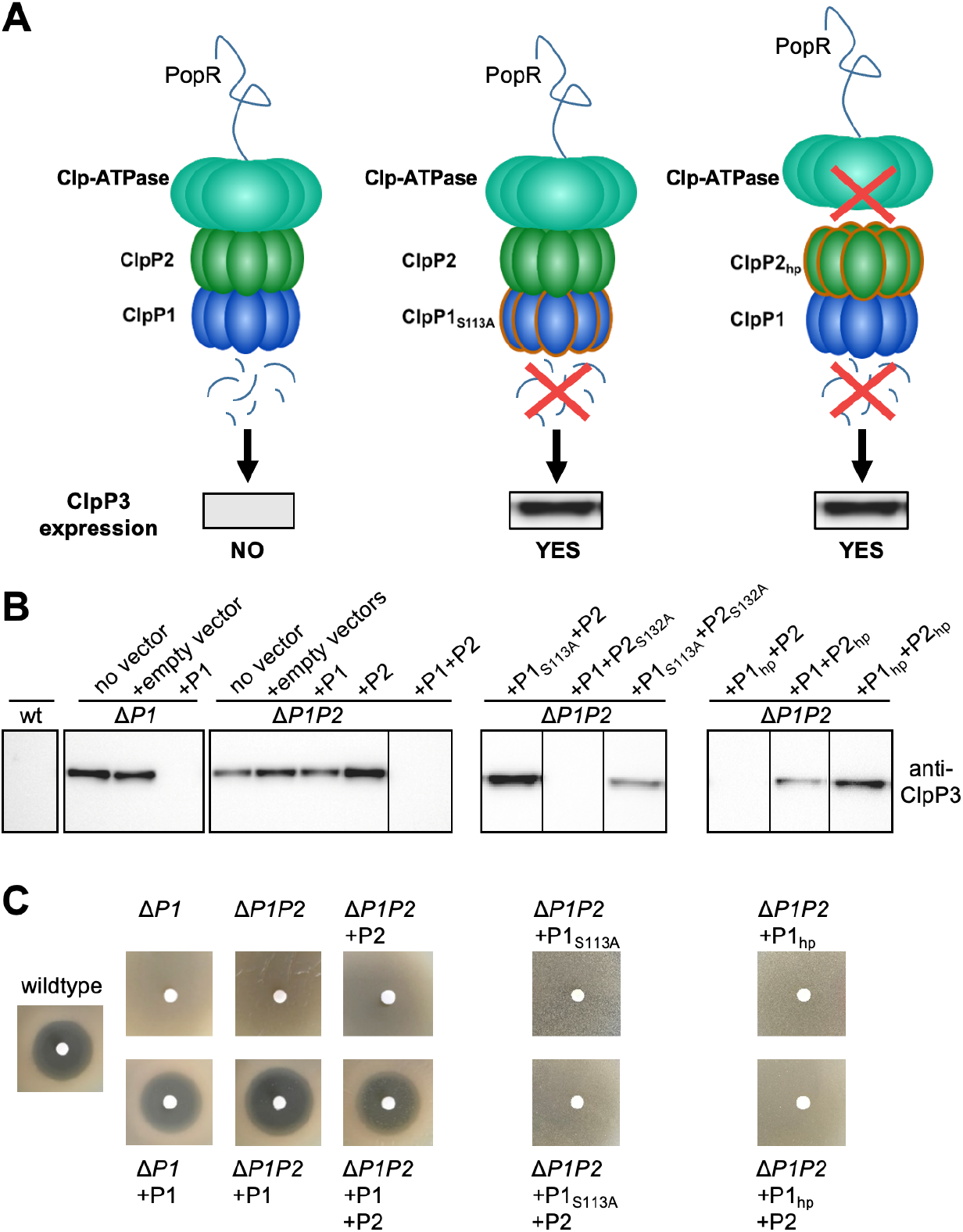
Whole cell studies verify the presence of a heteromeric ClpP1P2 complex, Clp-ATPase binding via ClpP2 as well as ADEP binding via ClpP1. **A.** Schematic overview of the effect of Clp protease functionalities on ClpP3 expression. **B.** Immunoblotting of cell lysates from wild-type and mutant *S. lividans* cells (as indicated) using anti-ClpP3 antibodies. By reconstituting wild-type, catalytic triad mutants, or hydrophobic pocket mutants of SlClpP1 or SlClpP2, whole cell assays underline the requirement for both SlClpP1 and SlClpP2 to build a functional, heteromeric Clp protease system, which strictly depends on a functional catalytic triad of SlClpP1 for proteolytic activity and on a functional hydrophobic pocket of SlClpP2 for the interaction with corresponding Clp-ATPases (both indicated by the loss of PopR degradation and concomitant expression of SlClpP3 in mutants harbouring either ClpP1_S113A_ or ClpP2_hp_, respectively). All assays were performed at least in triplicates. Representative Western Blot images are shown. **C.** Disk diffusion bioassay using ADEP1 verifies ADEP binding via ClpP1 in *S. lividans*, since only the loss of a functional copy of SlClpP1, but not of SlClpP2, confers ADEP resistance. Bioassays were performed at least in triplicates. Representative images are shown.

We then explored ClpP catalytic triad mutant proteins in *S. lividans* by complementing the Δ*SlP1P2* mutant with different combinations of the wild-type and serine-to-alanine mutant genes *SlclpP1_S113A_, SlclpP2_S132A_*, and *SlclpP2_S132A_-His* (Fig 4B, S5B). Here, immunoblotting experiments showed that a SlClpP1P2 system capable of SlPopR degradation depends on a functional catalytic triad in SlClpP1, but not in SlClpP2, thus supporting our *in vitro* results and providing first *in vivo* evidence that the catalytic function of ClpP1 accounts for the proteolytic activity of the heteromeric ClpP1P2 complex in *Streptomyces*.

We next investigated the binding of Clp-ATPases to the SlClpP1P2 complex in *S. lividans* cells by complementing the Δ*SlP1P2* mutant with different combinations of the wild-type and hydrophobic pocket mutant proteins SlClpP1_hp_, SlClpP2_hp_, and SlClpP2_hp_-His (Fig 4B, S5C). The degradation of SlPopR and thus the presence of a SlClpP1P2 system capable of hydrolysing protein substrates, depended on a functional hydrophobic pocket in SlClpP2, but not SlClpP1, therefore corroborating our *in vitro* results that Clp-ATPase binding occurs via ClpP2, but not via ClpP1. Regarding the binding of ADEP to the SlClpP1P2 complex, we conducted disk diffusion bioassays to determine the effect of the respective deletions and complementations on ADEP sensitivity (Fig 4C). Here, ADEP sensitivity entirely relied on the presence of a functional SlClpP1 protein, but not on SlClpP2, thus verifying the exclusive binding of ADEP to ClpP1 and indicating that a functional ClpP1 protein is sufficient to unleash the antimicrobial activity of ADEP in *Streptomyces*.

### ADEP antibiotics accelerate the Clp-ATPase-dependent hydrolysis of protein substrates by the *Streptomyces* Clp protease

Our data hitherto described provide *in vitro* and *in vivo* evidence that *Streptomyces* ClpP1 and ClpP2 form a hetero-tetradecameric complex that can be activated for proteolytic activity either by a cognate Clp-ATPase via ClpP2 or by the small molecule antibiotic ADEP1 via ClpP1. In previous studies reporting on Clp proteases from other bacteria such as *B. subtilis, Staphylococcus aureus, E. coli, M. tuberculosis* and *Chlamydia trachomatis*, the addition of ADEP to a reaction mixture of ClpP and a Clp-ATPase abrogated the interaction between the Clp-ATPase and the ClpP core in all cases and consequently prevented Clp-ATPase-dependent substrate hydrolysis without exception (19, 37–40). This steric competition led to the inhibition of the natural functions of the Clp protease in all of these bacteria. Intrigued by our observation that the *Streptomyces* Clp-ATPases and ADEP interact with separate ClpPs, we set out to explore if the situation might be different in this genus. First, we compared *in vitro* protein degradation of the native substrates ClgR and PopR by ClpXP1P2 in the absence or presence of ADEP1. While ClgR and PopR were digested by ClpXP1P2 in the absence of ADEP (Fig 1B, 2D), the addition of ADEP did not prevent the degradation of both substrates but, on the contrary, even led to a weak but notable increase of substrate digestion over time (Fig 5AE). Also, the ClpC1P1P2-mediated degradation of PopR was markedly stimulated by ADEP (Fig S6A). Of note, protein degradation did not occur in the absence of ClpX or with the hydrophobic pocket mutant protein ClpP2_hp_ (Fig 5AE), indicating that the interaction with a Clp-ATPase remains a prerequisite for the digestion of natural Clp substrates and excluding a Clp-ATPase-independent digestion of both substrates via ADEP-activated ClpP1P2.

**Figure 5:**
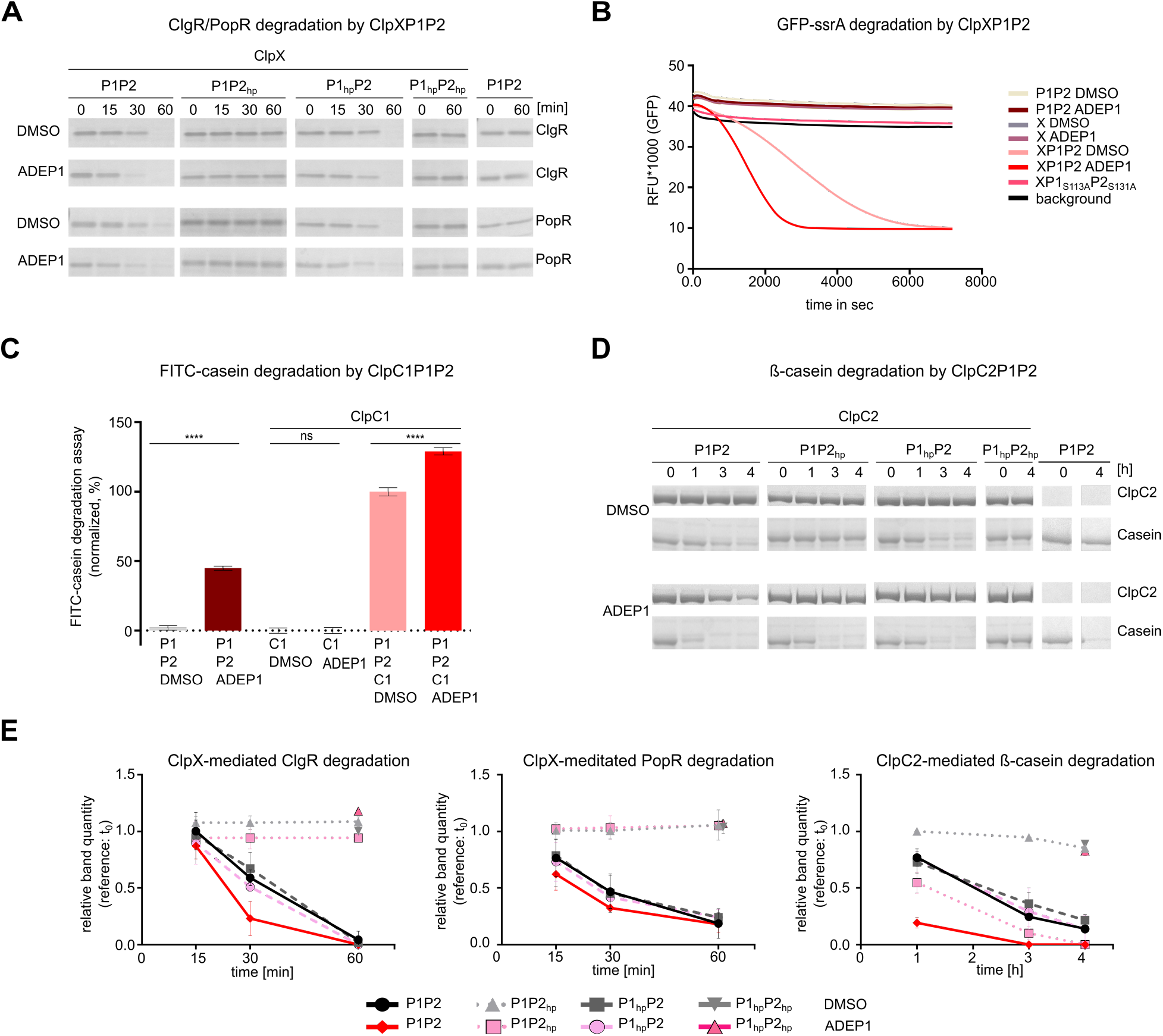
ADEP1 accelerates the proteolytic activity of the *Streptomyces* Clp protease. **A.** ADEP accelerates the degradation of the natural Clp substrates ClgR and PopR by ClpXP1P2 in *in vitro* protein degradation assays. Functional ClpX and its interaction with ClpP2 are prerequisites for ClgR and PopR degradation, even in the presence of ADEP. **B.** ADEP increases the hydrolysis of GFP-ssrA by ClpXP1P2. GFP-ssrA degradation was measured via fluorescence decrease over time. Background indicates default GFP fluorescence in the buffer solution without enzymes. Data shown are exemplary for at least three biological replicates. **C.** ADEP activates the free ClpP1P2 core to degrade FITC-casein and in addition stimulates the ClpC1-mediated FITC-casein degradation. Protease activity assays using FITC-casein as a substrate show that ADEP1 induces FITC-casein degradation by both ClpP1P2 as well as by ClpC1P1P2. Hydrolysis of FITC-casein was monitored as RFU increase over time. Mean values (normalized in %) of initial linear reaction kinetics are shown. P-values were calculated with one-way ANOVA from three biological replicates each comprising three technical replicates. P-values: ns > 0.05; **** ≤ 0.0001. Error bars indicate standard deviations. **D.** ADEP activates the free ClpP1P2 core to degrade ß-casein and in addition stimulates the ClpC2-mediated ß-casein degradation. ClpC2-mediated ß-casein degradation is prevented in the hydrophobic pocket mutant protein ClpP2_hp_. However, ADEP1 can still activate the ClpP1P2_hp_ proteolytic core by binding to ClpP1. Hydrolysis of ß-casein is only completely prevented when both hydrophobic pocket mutants, ClpP1_hp_ and ClpP2_hp_, are used in combination. Hydrolysis of ß-casein is fastest when wild-type ClpP1P2 interacts with both activators, i.e., ClpC2 plus ADEP, in parallel. Of note, in ADEP-containing reaction mixtures, ClpC2P1P2 degrades ß-casein first followed by digestion of ClpC2. All assays were performed at least in triplicates. In SDS-PAGE images, representative experiments are shown. DMSO was used in all control reactions. **E.** Densitometry of SDS-PAGE proteins bands of ClgR and PopR (Fig 5a) and ß-casein (Fig 5d) of each time point relative to the samples collected at t_0_. The relative band quantity was measured from three replicate SDS-PAGE analyses. Mean values are shown and error bars indicate corresponding standard deviations. Of note, a slight decrease of the ß-casein band in the negative control ClpP1_hp_P2_hp_ and ClpP1P2_hp_ could be detected, which might be due to independent unfolding activity of the only partially folded ß-casein by ClpC2.

To further study the unprecedented acceleration of the Clp-ATPase-dependent substrate hydrolysis by ADEP, we tested the effect of ADEP1 on the degradation of the protein model substrate GFP-ssrA by ClpXP1P2. The ssrA-tag is a common degron that is recognized by Clp-ATPases and results in the subsequent degradation of the fusion protein by the Clp protease (41). The kinetics of GFP-ssrA hydrolysis can be conveniently followed by a decrease in fluorescence (42). In line with our previous results, ADEP1 clearly stimulated GFP-ssrA degradation by ClpXP1P2 (Fig 5B). Neither ClpP1P2 alone nor ClpX alone, in the absence or presence of ADEP1, led to substrate degradation, demonstrating the strict necessity of a fully assembled Clp protease complex for GFP-ssrA hydrolysis. In our control sample with ClpXP1_S113A_ClpP2_S131A_, we did not detect a decline in RFU, indicating that the drop in RFU, measured for ClpXP1P2, is due to unfolding and cleavage of the substrate. Similarly, ADEP1 notably induced the degradation of FITC-casein by ClpC1P1P2 (Fig 5C) and ß-casein by ClpC2P1P2 (Fig 5D). Drawing this conclusion from experiments with casein, which can serve as a substrate for ADEP-activated as well as Clp-ATPase-activated ClpP1P2, is possible by comparing the degradation rates of the individual complexes. Degradation efficacy was the highest, when both activators were present at the same time (forming ADEP-ClpP1P2-Clp-ATPase complexes) and significantly lower (p ≤ 0.0001) for ClpP1P2-Clp-ATPase, ADEP-ClpP1P2 and ADEP-ClpP1 (activity decreasing in that order; Fig.3F, Fig 5C-E). The fact that ADEP addition to the ClpP1P2-Clp-ATPase complex further stimulated casein proteolysis speaks against significant formation of the less active ADEP-only complexes. Further noteworthy, the stimulated Clp-ATPase/ClpP1P2 activity in the presence of ADEP was not due to an increased ATP turn-over, since the conversion of ATP to ADP was unchanged in ClpC1-mediated casein degradation assays in the absence and presence of ADEP (Fig S6B). In summary, our data clearly show that ADEP1 accelerates the Clp-ATPase-driven degradation of protein substrates by the *Streptomyces* house-keeping Clp protease.

Further noteworthy, under these conditions and after ß-casein had been fully digested, ClpC2 was slowly degraded as well (Fig 5D). Similarly, after complete degradation of PopR had occurred and during prolonged incubation, also ClpC1 was degraded in the presence of ADEP1 (Fig S6A), suggesting ClpC1 and ClpC2 as putative targets of the ADEP-deregulated ClpP protease in *Streptomyces*.

## Discussion

Soil-dwelling bacteria of the genus *Streptomyces* are among the most important producers of secondary metabolites, facilitating the production of over two-thirds of all antibiotics in clinical use today (43). Such vast biosynthetic capacity of streptomycetes, which is accompanied by a complex developmental life cycle, clearly requires tight regulation and coordination of the involved processes. Regulated proteolysis has emerged as an important level of regulation that is intricately linked to cell-cycle progression and physiological transitions (44). We here set out to study the molecular function of the house-keeping *Streptomyces* Clp protease machinery, an essential compartmentalized protease in *Streptomyces* and a major player of regulated proteolysis in bacteria (3–5, 45). The *Streptomyces* Clp system is one of the most complex known to date, since streptomycetes encode up to five different ClpP homologs and at least four different Clp-ATPases (3, 21). Earlier *S. lividans* whole cell studies by Mazodier and colleagues provided first important insights into the potential working mode of the *Streptomyces* Clp protease. They observed that the presence of ClpP1 and ClpP2 precluded the expression of ClpP3 and ClpP4, and that the presence of both ClpP1 and ClpP2 was necessary for the digestion of PopR or ClgR (22, 23). Hence, their data suggested the presence of house-keeping Clp proteases with one or more proteolytic cores, consisting of either ClpP1, ClpP2 or even both homologs, as well as a back-up conferred by ClpP3 and/or ClpP4 that takes over the most important cellular functions in case of a dysfunction of the house-keeping Clp machinery.

Almost two decades later, we have now deciphered the composition and molecular operation mode of the house-keeping Clp protease in *Streptomyces* via *in vitro* reconstitution of the protease system side-by-side with corresponding whole cells studies. We here prove that the *Streptomyces* house-keeping Clp protease contains a hetero-tetradecameric, proteolytic core of both ClpP1 and ClpP2 that interacts with the corresponding Clp-ATPases ClpX, ClpC1 or ClpC2 for the degradation of proteins, such as the natural Clp substrates ClgR and PopR. Within the assembled Clp protease variants, we show that ClpP1 and ClpP2 fulfil distinct but complementary functions with regard to substrate hydrolysis. Our results clearly indicate that mainly ClpP1 confers catalytic activity to the proteolytic core, while ClpP2 is essential for the interaction with the corresponding Clp-ATPases to allow for substrate unfolding and translocation into the degradation chamber of ClpP1P2.

Our data further indicate that the heteromeric ClpP1P2 core is formed by two separate homo-heptameric rings of either ClpP1 or ClpP2, since functional Clp-ATPases are hexamers and all six IGF-loops of *E. coli* ClpX (EcClpX) are required for productive substrate digestion by EcClpXP (46). In that study, even the loss of a single IGF-loop reduced the affinity of EcClpX to EcClpP by approximately 50-fold. However, in the *Streptomyces* ClpP1P2 core, mutating the hydrophobic pocket of ClpP1 did not at all affect the hydrolysis of native substrates, which clearly argues against the presence of even a single ClpP1 monomer in the Clp-ATPase-interacting ClpP2 ring. Also, both *Streptomyces* ClpP1 as well as ClpP2 exist as homo-heptamers as revealed by size exclusion chromatography in our study, making the presence of mixed heptamers in the active protease highly unlikely. The presence of two homo-heptameric rings of either ClpP1 or ClpP2 in *Streptomyces* further correlates with the composition of the ClpP system in the closely related *M. tuberculosis* (33, 47, 48), *Mycobacterium smegmatis* (20), and *L. monocytogenes* (16, 49), which form hetero-tetradecamers composed of a ClpP1 heptamer stacked to a ClpP2 heptamer and those complexes bind Clp-ATPases solely via ClpP2.

*S. hawaiiensis* is the producer of the ClpP-interfering antibiotic ADEP1, the natural product progenitor of a promising class of potent acyldepsipeptide antibiotics (26, 28, 31, 50). But although the ADEP mode of action has been investigated in more detail in the major target bacteria, the effects of ADEP on the house-keeping Clp protease of the producer genus *Streptomyces* have remained largely elusive, so far. Pioneering whole cell studies by Mazodier and colleagues using the ADEP-sensitive *S. lividans* showed that ClpP1 is a target for ADEP1, since the deletion of the *clpP1* gene conferred ADEP resistance in *S. lividans* (36). In addition, we have very recently reported on the presence of a self-resistance factor in the ADEP producer *S. hawaiiensis* NRRL 15010, which is an accessory ClpP protein with a yet unknown mechanism (26). To obtain further insights into the role of ADEP in *Streptomyces*, we here investigated the effect of ADEP on the molecular function of the house-keeping Clp protease systems of *S. hawaiiensis* and *S. lividans*. Of note, the ClpP1 and ClpP2 proteins of *S. hawaiiensis* show very high amino acid sequence similarities to their corresponding homologs in *S. lividans*, thus suggesting a similar operation mode of ClpP1P2 in both species (Fig S1D).

In previous studies on the antibacterial mode of action of ADEP antibiotics, two distinct molecular mechanisms were identified that, depending on the bacterial species and growth environment, together or individually lead to bacterial killing. First, since Clp-ATPases and ADEP accommodate the same binding site on ClpP, binding of ADEP to ClpP displaces the commonly associated Clp-ATPases from the proteolytic core (19, 37–40), which leads to an inhibition of all physiological functions of the Clp protease in the bacterial cell including regulated proteolysis. Second, ADEP binding induces an opening of the ClpP entrance pores and allosterically activates the catalytic centers, thereby allowing access of non-native substrates to the ClpP degradation chamber that are efficiently degraded (19, 30, 34, 35, 38, 51, 52). The latter activation of the independent proteolytic core ClpP is the primary cause of bacterial killing in species harbouring non-essential ClpP proteins, such as *B. subtilis* or *S. aureus* (27, 29, 30, 51, 53, 54), although in the infection situation, where the Clp protease is essential for the expression important virulence factors, inhibition of the natural functions will also attenuate a pathogen in the host environment (55, 56). In *M. tuberculosis*, the Clp protease is essential for viability under all conditions (48, 57), and we have previously shown that an ADEP-conferred mechanism of Clp protease inhibition leads to cell death in mycobacteria (39). Considering the essentiality of the Clp protease in *Streptomyces* and the comparative phylogenetic closeness to mycobacteria (21), it was thus tempting to hypothesize that a mechanism of Clp protease inhibition could be the basis of ADEP-dependent killing of *Streptomyces*. However, to our surprise, ADEP did not abrogate the Clp-ATPase-mediated substrate digestion. Actually, all of our data consistently show that the presence of ADEP leads to an unexpected stimulation of the *Streptomyces* Clp protease to digest its natural substrates in a Clp-ATPase-dependent, more efficient manner, rather than leading to an inhibition of its natural functions. Furthermore, ADEP-stimulated Clp-ATPase/ClpP1P2 activity was not due to an increased ATP turn-over, hence, stimulation may be solely attributed to the ADEP effects that are exerted on the structural network of the entire ClpP1P2 complex, which may include adjustment of the active site residues in ClpP1 as well as pore opening for ClpP1 and ClpP2, as was previously described for ADEP-activated ClpP in other bacteria (15, 34, 35, 38). Therefore, the current study reveals a third, yet unobserved mode of ADEP action.

This new ADEP effect observed here can be explained by our observation that, in *Streptomyces*, the proteolytic ClpP1P2 core can be powered simultaneously by two different activator types. We found ADEP to bind solely to ClpP1 of *S. hawaiiensis* and *S. lividans*. Of note, very recently, asymmetric binding of ADEP to ClpP1 was also observed for ClpP1P2 hetero-tetradecamers of *Streptomyces cattleya* (58) suggesting that this binding mode is a general characteristic of the genus *Streptomyces*. Furthermore, our data consistently show, that the ClpP2 apical face of the *Streptomyces* ClpP1P2 core serves as the sole docking site for three different Clp-ATPases, a fact that was elusive before, as the *S. cattleya* study did not include work on Clp-ATPases. The simultaneous and unhindered binding of ADEP and a Clp-ATPase to opposite sides of the *Streptomyces* ClpP core is unique among all the Clp systems investigated so far.

Considering previous studies on ClpP proteins from other bacteria and the effects of ADEP antibiotics, structural analyses identified three different conformations of the ClpP tetradecamer, a proteolytically active extended conformation, an inactive compact intermediate and an inactive compressed conformation, which all differ in the height of the ClpP barrel as well as regarding the alignment of the catalytic triad residues (14, 59–63). The Clp protease core is understood to dynamically switch between these different conformations, thereby exhibiting separate steps of substrate hydrolysis and product release during processive substrate degradation (49, 64, 65). Here, degradation products might exit the degradation chamber through transient equatorial pores that emerge in the proteolytically inactive compressed conformation rather than through the axial pores that may be blocked by the partner Clp-ATPases (62). Importantly, we and others have previously shown that the binding of ADEP to ClpP leads to long-distance conformational changes, thereby opening the entrance pores to the degradation chamber and locking the ClpP tetradecamer in the active extended conformation (15, 34, 35, 38). In our current study, binding of ADEP1 led to the oligomerization of *Streptomyces* ClpP1 into proteolytically active ClpP1 homo-tetradecamers and stimulated proteolysis by ClpP1P2_hp_ hetero-tetradecamers carrying a defective ClpP2 hydrophobic pocket. Both findings clearly indicate the capacity of ADEP1 to open the entrance pores to the degradation chamber by docking to *Streptomyces* ClpP1, since this is a prerequisite for the entry of protein substrates and subsequent protein digestion (9, 10). We therefore propose the following model of ADEP action in *Streptomyces* (Fig 6): the partner Clp-ATPase binds to the ClpP1P2 proteolytic core via ClpP2 and unfolds and feeds the natural protein substrates into the degradation chamber. The simultaneous binding of ADEP to the opposite side of the proteolytic core via ClpP1 stabilises the core in the proteolytically active extended conformation (including the correct arrangement of the active centres for catalysis) and stimulates the proteolysis of natural Clp substrates that are fed by the Clp-ATPase via ClpP2. This process may be further accelerated by a more efficient product release through an opened pore of the ADEP-bound ClpP1 heptamer of the proteolytically-active tetradecameric core. We have previously shown that ADEP-activated EcClpP degrades largely unstructured proteins like casein with reduced processivity compared to EcClpAP (37). This may be explained by the diffusion of partially cleaved substrates through the opened axial pores in ADEP-activated EcClpP, thereby impeding processive cleavage, whereas such escape of degradation intermediates is prevented in EcClpAP (35). In our model for the *Streptomyces* Clp protease, however, Clp-ATPase-dependent processive cleavage would be allowed even in the presence of ADEP, potentially stimulating the degradation of natural Clp substrates. In the closely related *M. tuberculosis*, ADEP binds to only one side of the ClpP hetero-tetradecamer exclusively (i.e., to MtClpP2) but was shown in crystal structures to open the entrance pores on both sides of the barrel (15). Thus, it may be further hypothesized that upon binding of ADEP to *Streptomyces* ClpP1, long-distance conformational shifts might also widen the diameter of the ClpP2 axial pore, which could additionally ease the feeding of proteins substrates by the Clp-ATPase.

**Figure 6:**
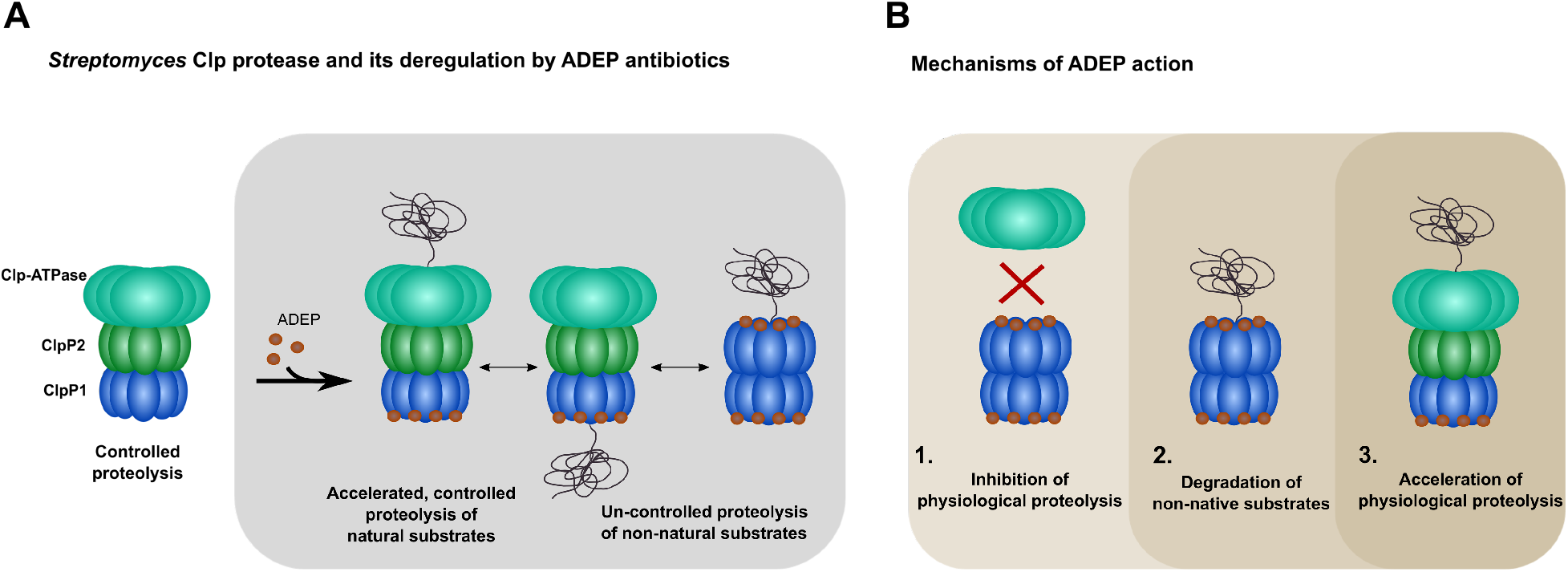
Mechanisms of action of ADEP antibiotics in *Streptomyces* and other bacteria. **A.** Model of the house-keeping Clp protease in *Streptomyces* and its deregulation by ADEP antibiotics. The major Clp protease in *Streptomyces* is composed of a hetero-tetradecameric core formed by ClpP1 and ClpP2 that interacts with the AAA+ chaperones ClpX, ClpC1 or ClpC2 via ClpP2. The catalytic triad of ClpP1 is solely responsible for enzymatic hydrolysis of protein and polypeptide substrates. Binding to the ClpP1P2 proteolytic core via ClpP2, the Clp-ATPases unfold and feed natural protein substrates into the degradation chamber in an ATP-dependent manner. Simultaneous binding of ADEP can occur at the opposite side of the proteolytic core via ClpP1, which locks the core in the proteolytically active extended conformation and stimulates the proteolysis of natural Clp substrates that are fed by a Clp-ATPase via ClpP2. Binding of the antibiotic to ClpP1 also widens the ClpP1 entrance pore to allow entry of non-native substrates into the ClpP1P2 hetero-tetradecamer. In addition, ADEP triggers the oligomerization of ClpP1 into a homo-tetradecamer, which is thereby activated for Clp-ATPase-independent proteolysis of non-native substrates. As the presence of ClpP1 alone (i.e., in the absence of ClpP2) is sufficient to sensitize *Streptomyces* against ADEP, the degradation of non-native protein and polypeptide substrates is the primary reason for cell death in this species. Nonetheless, while the physiological protein substrates of the Clp protease can still be degraded in the presence of ADEP, accelerated degradation of native substrates will probably also stress the cells through protein imbalance. **B**. Summary of mechanisms of action of ADEP antibiotics. We have previously described that ADEP antibiotics can lead to the inhibition of natural Clp protease functions (1) as well as to the uncontrolled degradation on non-natural Clp substrates in the absence of otherwise regulating Clp-ATPases (2). We here present a third, new mechanism of ADEP action, that manifests in the producer genus *Streptomyces* that leads to the acceleration of the Clp-ATPase-dependent digestion of natural Clp protease substrates (3).

Indeed, our results clearly indicate the existence of an inherent communication network between the ClpP1 and ClpP2 heptameric rings of the *Streptomyces* ClpP barrel. For example, the binding of a Clp-ATPase to ClpP2 led to catalytic stimulation of ClpP1, potentially implying an allosteric activation mechanism for ClpP1 upon concerted conformational rearrangements in both heptameric rings. Also, self-processing of ClpP1 was induced by the binding of ClpX to ClpP2 at the opposite side of the heteromeric complex *in vitro*. Inter-ring communication is not unusual among Clp proteases studied to date (66–69), for example, Weber-Ban and colleagues showed that in asymmetric *E. coli* ClpP tetradecamers composed of one wild-type and one catalytically inactive heptameric ring, ClpA-ATPase activity could be stimulated by ClpP regardless of whether ClpA bound to the wild-type or the catalytically inactive ring (70). Previously it has also been shown that ClpA activity could only be activated in the presence of catalytically active ClpP proteins by an allosteric signal across the ClpA-ClpP interaction surface (71).

The results presented in this study unravel the composition and molecular operation mode of the *Streptomyces* house-keeping Clp protease machinery as well as its unique stimulation by ADEP antibiotics. Although the protein imbalance upon enhanced degradation of natural Clp substrates can be expected to cause severe cellular stress, the ADEP-mediated catalytic acceleration of the Clp-ATPase-controlled digestion of natural Clp substrates is definitely not the only kind of deregulation that ADEP inflicts on the *Streptomyces* Clp protease machinery. The efficient degradation of casein by ADEP-bound ClpP1P2 and ClpP1 shows that non-native substrates gain access to both ADEP-activated proteolytic cores, i.e., the native ClpP1P2 hetero-tetradecamer as well as the presumably non-native ClpP1 homo-tetradecamer, via the opened entrance pore of the ClpP1 heptameric ring. The fact that *S. lividans* mutants carrying ClpP1 but lacking ClpP2 were killed by ADEP application illustrates that Clp-ATPase-independent and thus uncontrolled proteolysis events of essential and non-essential polypeptide or protein substrates are already sufficient to cause cell death in *Streptomyces*.

## Methods

### Construction of plasmids for protein expression

Plasmids for the expression of *clp, clgR* and *popR* genes of *S. hawaiiensis* NRRL 15010 were obtained by ligating PCR-amplified gene sequences with linearized pET11a or pET22b vectors (Merck-Novagen), respectively. All bacterial strains, plasmids and primers used in this study are listed in tables S1 and S2, including a detailed description of templates and restriction enzymes used. As PCR template, genomic DNA of *S. hawaiiensis* NRRL 15010 (NCBI Genbank accession number: CP021978.1) was used to amplify *shclpP1* (orf CEB94_14110), *shclpP2* (orf CEB94_14105), *shClpX* (orf CEB94_14100), *shclpC1* (orf CEB94_23085), *shclpC2* (orf CEB94_33910), *shclgR* (orf CEB94_30145), and *shpopR* (MT943519). Since *clpP2* encodes an internal *Xho*I restriction site, the vector pET22b\**Nco*I was created, in which the *Xho*I restriction site was exchanged for an *Nco*I restriction site (19). In addition, both *shclpP1* and *shclpP2* genes were cloned into the pETDUET™-1 vector (Merck-Novagen), yielding the plasmid *pETDUETshclpP1_ATG2_clpP2-His*, in which the simultaneous expression of the *clp* genes is regulated by two distinct T7 promoters. Additionally, side-directed mutagenesis was performed, yielding the constructs pET22bshclpP1_S113A_, pET22bshclpP2_S131A_, pET11ashclpP1_hp_, pET11ashclpP1_Y76S_, pET22bshclpP2_hp_, pET22bshClpP2_S94Y_. Therefore, primers were designed according to the protocol of the QuickChange II Site-directed mutagenesis kit (Agilent) and as listed in table S2. To verify the correct nucleotide sequence of the respective gene, Sanger sequencing was performed by LGC genomics, Germany. During sequence analysis of ClpP2, a second putative start codon was detected, which would give rise to a shortened protein (ClpP2_ATG2_). To ensure, that ClpP2_ATG2_ would behave similar compared to full-length ClpP2, we also cloned and purified ClpP2_ATG2_, determined its processing site, and performed degradation experiments using ClpP2 and ClpP2_ATG2_ side by side. Here, no differences were observed for ClpP2_ATG2_ compared to using ClpP2 (Fig S7). Of note, due to solubility issues, ClpP2 proteins with mutations in the hydrophobic pocket (i.e., ClpP2_hp_, ClpP2_S94Y_) were produced from ATG2, and were then compared to ClpP2_ATG2_ in respective assays (Fig 2B-D, 3B, 3E, 5AD, S2G, S3A, S7).

### Protein expression and purification

For the expression and purification of Clp proteins and the substrates ClgR and PopR, the respective expression plasmids were transformed into the Δ*clpP* deletion strain *E. coli* SG1146a (72). Resulting expression strains were then used to inoculate expression cultures using lysogeny broth (LB) supplemented with ampicillin (100 μg/ml). Cultures were grown at 37 °C with constant shaking until a mid-log exponential growth-phase (OD_600_ of 0.5 - 0.6) was reached and then protein expression was induced with 0.5 - 1 mM IPTG. Expression cultures for ClpP1, the ClpATPases and PopR were further incubated for 4 - 5 h at 30 °C with constant shaking, whereas ClpP2 and ClgR expression was carried out for 16 h at 30 °C and 20 °C, respectively. Then, cells were harvested by centrifugation. The following purification procedure was performed at 4 °C. For cell disruption, glass beads were utilized (150-212 μm) in a Precellys homogenizer (Precellys evolution, Bertin technologies). Cell lysates were centrifuged, and supernatants were passed through 0.45 μm membrane filters (Sarstedt) to remove cell debris before further processing. For purification of untagged ClpP1 and ClpP2, the respective supernatants were applied to 1/5 ml HiTrap™ Q XL columns (GE Healthcare) for anion exchange chromatography using the ÄKTA Start chromatography system (GE Healthcare) with buffer A (50 mM Tris, pH 8) and buffer B (50 mM Tris, 1 M NaCl, pH 8). For purification of His_6_-tagged proteins, 500 - 700 μl Ni-NTA resin (Thermo-Fisher) was added to the supernatant and the suspension was incubated for at least 2 h at 4 °C. Affinity chromatography was performed using lysis buffer (50 mM NaH_2_PO_4_, 300 mM NaCl, 10 mM imidazole pH 7.7), wash buffer (50 mM NaH_2_PO_4_, 300 mM NaCl, 20 mM imidazole, pH 7.7), and elution buffer (50 mM NaH_2_PO_4_, 300 mM NaCl, 500 mM imidazole pH 7.7). Protein fractions were pooled for buffer exchange employing a PD-10 Sephadex™ G-25 desalting column (GE Healthcare) and centrifugal filters (Amicon®Ultracel®-10 K/30 K; Merck). Protein concentration and purity was determined via SDS PAGE analysis, Bradford assay (Bio-Rad, applying BSA as reference), and immunoblotting. Here, polyclonal rabbit anti-ClpP1 and rabbit anti-ClpP3 antibodies (provided by Dr. Philippe Mazodier) were used, which had been previously described (22-24, 36). For 6-His-tag detection, monoclonal mouse anti-6x-His antibodies (Epitope-tag-clone: HIS.H8, Thermo Fisher) were employed. Accordingly, IgG (H+L) rabbit anti-mouse HRP (Invitrogen) or IgG (H+L) goat anti-rabbit HRP (Pierce) were used as secondary antibodies.

### Size exclusion chromatography

To study the oligomeric behaviour of ClpP1 and ClpP2, analytical gel filtration was performed using the Superdex™ 200 Increase 3.2/300 column (GE Healthcare). To this end, 80 μM of each protein were incubated with or without the addition of 200 μM ADEP1 for 30 min at 30 °C. As a control, DMSO was used instead of ADEP1. Additionally, ClpP1P2_His_ (90 μM each) were incubated with 200 μM ADEP1 for 2 h at 30 °C and applied to the gel filtration column. For gel filtration, buffer GF was used (50 mM HEPES-KOH, 150 mM KCl, 20 mM MgCl2, 10% glycerol, pH 7.5). The protein markers thyroglobulin (669 kDa), apoferritin (443 kDa), ß-amylase (200 kDa), alcohol dehydrogenase (150 kDa, yeast), BSA (66 kDa) and carbonic anhydrase (29 kDa, bovine) were employed for size estimation.

### *In vitro* peptidase assays

Peptidase assays were carried out in a total volume of 100 μl at 30 °C in activity buffer (50 mM HEPES-KOH, 150 mM KCl, 20 mM MgCl2, 10% glycerol, pH 7.5). Peptidase activity of ClpP1 and ClpP2_His_ was measured using 100 μM fluorogenic N-succinyl-Leu-Tyr-7-amido-4-methylcoumarin (Suc-LY-AMC), 100 μM benzyloxycarbonyl-glycyl-glycyl-L-leucine-7-amino-4-methylcoumarin (Z-GGL-AMC), 100 μM acetyl-Trp-Leu-Ala-7-amino-4-methylcoumarin (Ac-WLA-AMC) or 100 μM Ac-Ala-hArg-(*S*)-2-aminooctanoic acid-7-amino-4-carbamoylmethylcoumarin (Ac-Ala-hArg-2-Aoc-ACC) as substrates in the absence or presence of 10 μM ADEP1. A total ClpP protein concentration of 4 μM was employed for each sample, using equal amounts of ClpP1 and ClpP2 (2 μM each) in mixed samples. Furthermore, ClpP2 (4 μM) was tested at increased concentrations of ADEP1, i.e., 100 μM and 200 μM, while ADEP-sensitive ClpP1 (4 μM) plus 100 μM ADEP1 was used as a positive control. In all assays, DMSO served as a negative control. For the analysis of the hydrophobic pocket mutants ClpP1_hp_ and ClpP2_hp_, a total ClpP protein concentration of 8 μM in the absence or presence of 30 μM ADEP1, ADEP2 or ADEP4 were used. ClpP and ADEP/DMSO were pre-incubated for 10 min at 30 °C. The release of AMC (Suc-LY-AMC or Z-GGL-AMC) was monitored in black flat-bottom 96 well plates (Sarstedt) in a spectrofluorometer (TECAN Spark®) at an excitation wavelength λex of 380 nm and an emission wavelength λem of 460 nm. Experiments using Ac-WLA-AMC or Ac-Ala-hArg-2-Aoc-ACC as substrates were performed in black flat-bottom 96 well plates (Sarstedt) in a spectrofluorometer (TECAN Spark®) at λex 430 nm / λem 351 nm for Ac-WLA-AMC and λex 380 nm / λem 430 nm for Ac-Ala-hArg-2-Aoc-ACC. At least three biological replicates were analysed, each with three technical replicates, with an exception for testing ClpPhp mutants with two biological replicates. P-values were calculated with one-way ANOVA (P-values: ns > 0.05; * ≤ 0.05; ** ≤ 0.01; *** ≤ 0.001; **** ≤ 0.0001).

### In vitro casein degradation and ClpP processing assays

For the ClpC1-mediated degradation of fluorescein isothiocyanate-casein (FITC-casein; Sigma https://www.sigmaaldrich.com/DE/de/product/sigma/c3777), 4 μM ClpC1 and a total ClpP protein concentration of 8 μM (4 μM each in mixed samples) were resuspended in activity buffer. Proteins were pre-incubated for 10 min at 30 °C before the reaction was started by adding 6 μM FITC-casein. Mixed samples of ClpP1P2_His_ in the absence of ClpC1 were tested in control reactions. Reactions were performed in the absence or presence of 30 μM ADEP1 or equal volume of DMSO as a control. Degradation of FITC-casein (6 μM) by ClpC2P1P2 or ClpXP1P2 (4 μM each) was conducted accordingly. The hydrolysis of FITC-casein was monitored in black 96-well plates (Sarstedt) in a spectrofluorometer (Tecan Spark®) at λex 490 nm / λem 525 nm. At least with three biological replicates were analysed, each comprising three technical replicates. Statistics were performed with one-way ANOVA (P-values: ns > 0.05; * ≤ 0.05; ** ≤ 0.01; *** ≤ 0.001; **** ≤ 0.0001). For the ClpC2-mediated degradation of ß-casein, equal amounts of ClpP1, ClpP2_His_ and ClpC2 (1.5 μM each) and 10 μM ß-casein were incubated in activity buffer for 4 h at 30 °C. Reactions were performed in the absence or presence of 30 μM ADEP1 or equal volume of DMSO as a control. For *in vitro* ClpP processing assays, 2 μM of either ClpP1 or ClpP2_His_, or mixed samples consisting of both 2 μM ClpP1 and 2 μM ClpP2_His_ were incubated at 30 °C with 10 μM ADEP1 or DMSO in activity buffer. Samples were taken at indicated time points and were analysed via SDS-PAGE. Experiments were performed at least in biological triplicates and representative SDS-PAGE images are shown. In all assays for FITC-casein and ß-casein degradation, an artificial ATP regeneration system (4 mM ATP, 8 mM creatine phosphate, and 10 U /ml creatine phosphokinase) was used in the reactions to sufficiently replenish ATP for Clp-ATPase activity.

### *In vitro* degradation of ClgR and PopR

For the ClpX-mediated degradation of ClgR and PopR (3 μM each), 3 μM ClpP1, 3 μM ClpP2_His_ (or 1.5 μM each in mixed samples) were used in combination with 1.5 μM ClpX in activity buffer plus 2 mM DTT. For the ClpC1- or ClpC2-mediated degradation of ClgR and PopR (3 μM each), 1.5 μM ClpP1 and 1.5 μM ClpP2_His_ were used in combination with either ClpC1 or ClpC2 (each 1.5 μM). Reactions were performed in the absence or presence of 30 μM ADEP1 or an equal volume of DMSO as a control. Experiments were performed at least in biological triplicates and representative SDS-PAGE images are shown. An artificial ATP regeneration system (4 mM ATP, 8 mM creatine phosphate, and 10 U /ml creatine phosphokinase) was used in the reactions to sufficiently replenish ATP for Clp-ATPase activity.

### Densitometry

Relative quantity of protein bands was calculated using Image Lab Software (Biorad). As reference, the protein bands collected at time point 0 h (t0) were utilized for each sample. Mean values of three biological replicates are shown.

### *In vitro* degradation of GFP-ssrA

For the ClpX-mediated degradation of C-terminally ssrA-tagged GFP, 3 μM of ClpP1, ClpP2_His_ and ClpX were used, mixed in activity buffer plus 2 mM DTT. Reactions were performed in the absence or presence of 30 μM ADEP1 or equal volume of DMSO as a control. An artificial ATP regeneration system was added to the reactions as described above. All reaction components except for GFP-ssrA were pre-incubated for 10 min at 30 °C. Then, reactions were started by adding 0.36 μM GFP-ssrA. GFP fluorescence was monitored in white, flat bottom 96-well plates (Greiner) over 120 min at excitation wavelength λ ex of 465 nm and emission at λ em of 535 nm using a spectrofluorometer (TECAN infinite M200 and TECAN Spark®). The data shown is exemplary for three independently performed biological replicates.

### Pull-down assays

For the co-elution of untagged ClpP1 with His-tagged ClpP2, *E. coli* SG1146a harbouring the pETDUET-derived plasmid pETDUET*shclpP1clpP2*-His was cultured for 4 h at 30 °C before co-expression of both proteins was induced by the addition of 0.5 mM IPTG. Then, cells were harvested by centrifugation and lysed, and resulting supernatants were used for affinity chromatography as described above for His_6_-tagged proteins. The flow-through, wash and elution fractions were sampled and analysed via SDS-PAGE. As a control, purified ClpP1 or ClpP2_His_ (8 μM each) were passed in separate runs over a nickel-NTA chromatography column, and the flow-through, wash and elution fractions were collected and analysed. For immunoblotting, protein samples were transferred onto a nitrocellulose membrane (Thermo-Fisher) via semi-dry blotting for 1 h at 0.8 mA/cm^2^. For the detection of ClpP1 and ClpP2-His_6_, anti-ClpP1 (1:2000) and anti-His_6_ (1:1000) primary antibodies were employed. HRP-coupled secondary antibodies were used and resulting chemiluminescent signals were detected via a ChemiDoc documentation system (Biorad). Experiments were performed at least in biological triplicates and representative SDS-PAGE or Western blot images are shown.

### Intact-protein mass spectrometry

High-resolution intact protein mass spectrometry was performed to analyse the processing sites of ClpP1 and ClpP2_His_ of *S. hawaiiensis*. Measurements were performed on a Dionex Ultimate 3000 HPLC system coupled to an LTQ FT Ultra (Thermo) mass spectrometer with an electrospray ionization source (spray voltage 4.0 kV, tube lens 110 V, capillary voltage 48 V, sheath gas 60 a.u., aux gas 10 a.u., sweep gas 0.2 a.u.). Reaction mixtures containing a total of about 1 – 10 pmol protein were desalted with a Massprep desalting cartridge (Waters) before measurement. The mass spectrometer was operated in positive ion mode, collecting full scans at high resolution (R = 200,000) from m/z 600 to m/z 2000. The protein spectra were deconvoluted using the Thermo Xcalibur Xtract algorithm.

### Oligomerisation studies via native PAGE

In order to visualize the formation of a ClpP1P2 heteromeric complex, the cross-linking reagent BS3 (bis-sulfosuccinimidyl suberate; Thermo-Fisher) was used to stabilize interactions between ClpP1 and ClpP2_His_. To this end, ClpP1, ClpP2, or a mixture of both ClpP1 and ClpP2 (6 μM each) were pre-incubated for 30 min at 30 °C in activity buffer. For cross-linking reactions, a 50-fold molar excess of BS3 over the total ClpP concentration (6 mM BS3 for the mixed ClpP1P2 samples, or 3 mM BS3 for samples with either ClpP1 or ClpP2) was added to the samples. Reactions were incubated for 30 min at 30 °C without shaking. Subsequently, the reactions were stopped by adding 20 mM Tris to the samples followed by a further incubation for 15 min at room temperature. For analysing the oligomeric behaviour in the presence of ADEP1, ClpP1 and ClpP2 (10 μM each) were incubated with 60 μM ADEP1 (ClpP:ADEP1 ratio of 1:3) or 25 μM ADEP1 (ClpP:ADEP1 ratio of 1:1.25) for 3 h at 30 °C. DMSO was used as a negative control. Samples were subjected to native PAGE at 25 V for approx. 16 h at 4 °C. Gels were either Coomassie-stained or used for immunoblotting employing PVDF membranes (Merck) as described above. To visualize a heteromeric ClpP1P2 complex, cross-linking experiments were conducted using the following activity buffer: 50 mM HEPES, 200 mM Na_3_C_6_H_5_O_7_, 20 mM MgCl_2_, 10% glycerol, pH 7.5. The experiment was performed with ClpP1P2 (10 or 15 μM each), which were incubated for 30 min at 30 °C with ADEP1 (25 μM or 37.5 μM, respectively). Then, a 50-fold molar excess of BS3 over the total ClpP concentration was added to the samples and incubated for 30 min at 30 °C. Samples were analysed via native PAGE, using 4-12% Tris-glycine pre-cast gels and ready-mixed native sample buffer (both Thermo-Fisher). Immunoblotting was performed as described above except for using PVDF membranes.

### ATPase activity

The ADP-GloTM Kinase assay (Promega) was utilized to measure the turn-over of ATP to ADP in ClpC1-mediated casein degradation assays according to the manufacturer’s instructions. In a first step, the reaction was conducted using ClpP1P2_His_ (8 μM each), ClpC1 (8 μM), ß-casein (10 μM) as a substrate, as well as ultra-pure ATP (100 nM, Promega). After 60 min of incubation at 30 °C, the reaction was stopped by adding ADP-Glo^TM^ reagent to deplete remaining ATP. ATP depletion was carried out for 40 min at room temperature. In a second step, the Kinase Detection Reagent was added for the conversion of ADP to ATP and incubated for 1 h at RT using a 384-well plate. In 384-well plates, 5 μl reaction mixture, 5 μl ADP-Glo^TM^ reagent and 10 μl of Kinase Detection Reagent was used. The converted ATP was then measured via a luciferase/luciferin reaction and ADP-Glo^TM^ kinase assay luminescence was recorded.

### Bacterial strains, culture conditions and cloning procedures for whole cell studies

All bacterial strains, plasmids and primers used in this study are listed in tables S1 and S2. *Streptomyces* wild-type and mutant strains were grown at 30 °C on MS-MgCl2 agar (2% soy flour, 2% mannitol, 2% agar, 10 mM MgCl_2_) or in Tryptic Soy Broth (TSB; BD Biosciences) with apramycin (50 μg/ml) and/or hygromycin B (50 μg/ml) as appropriate. *E. coli* strains were propagated in LB medium with apramycin (50 μg/ml), hygromycin B (50 μg/ml), kanamycin (25 μg/ml), and/or chloramphenicol (30 μg/ml) as required. For cloning purposes, DNA fragments of the genes of interest were PCR-amplified and cloned into the respective vector via restriction-ligation utilizing *E. coli* JM109 or *E. coli* DH5α as cloning strains. All constructs were verified by Sanger sequencing (LGC Genomics). *Streptomyces* vectors were conjugated into the respective *Streptomyces* strains via *E. coli* ET12567 pUB307 as described previously (73). Site-directed mutagenesis was performed as described above. For the generation of *clpP* deletion mutants, markerless gene knockouts were achieved by PCR-amplifying 1.5 kb long flanking regions of the respective gene, which were then integrated into the pGM-GUS-Xba vector by Gibson assembly using the NEBuilder HiFi DNA Assembly Master Mix (NEB) and *E. coli* JM109 or E. coli DH5α as cloning hosts. After verification by Sanger sequencing, the knockout constructs were transformed into *E. coli* ET12567 pUB307 and conjugated into *S. lividans* TK24 as described previously (73). Mutant colonies were grown with apramycin at 39 °C in TSB liquid cultures and subsequently on MS agar to maintain cells with integrated plasmids only. Spores were harvested and dilutions were plated on LB agar with 2 μg/ml of ADEP1 as selective pressure for plasmid loss. The obtained colonies were overlaid with 1 ml of an X-Gluc (5-Bromo-4-chloro-3-indolyl-β-D-glucuronide) solution in water (1 mg/ml) for a blue-white screen. For white colonies, loss of the vector and the respective *clpP* gene were further verified by colony-PCR and subsequent Sanger sequencing.

### ADEP sensitivity assays

Spore suspensions of the respective *Streptomyces* strains were plated on Nutrient Extract (NE) agar (1% glucose, 0.2% yeast extract, 0.2% casamino acids, 0.1% Lab-Lemco Powder, pH 7.0). Paper disks containing 20 μg of ADEP1 were added to the plates, which were subsequently incubated at 30 °C for 2 days.

### Preparation of *Streptomyces* culture protein extracts

The respective *Streptomyces* strains were grown in 10 ml of TSB medium with apramycin and/or hygromycin as appropriate for 40 h at 30 °C and 180 rpm. The mycelium was harvested and resuspended in 500 μl of lysis buffer (20 mM Tris, 5 mM EDTA disodium salt, 1 mM 2-mercaptoethanol; one cOmplete™ Mini, EDTA-free protease inhibitor cocktail tablet (Roche) per 10 ml of lysis buffer) in 2 ml lysis tubes. Upon cell disruption using the Precellys homogenizer, cell debris was removed by two centrifugation steps (14,000 rpm, 10 min, 4 °C followed by 14,000 rpm, 20 mins, 4 °C) and the concentration of the protein extracts was determined by measuring the absorption at 280 nm with the NanoDrop 2000c spectrophotometer (Thermo Scientific).

## Acknowledgments and author contributions

We are grateful to all members of our teams and the RTG 1708 and TRR 261 for their support and critical discussions. Further, we thank Philippe Mazodier for the kind gift of antibodies, Günther Muth and Susan Gottesman for providing the bacterial strains *S. lividans* TK24 and *E. coli* SG1146a, respectively, Marc Buttner for sharing the plasmids pIJ12551 and pIJ10257, and we acknowledge the technical support of Stefan Pan, Janna Hauser and Denijel Latifovic.

L. Reinhardt, D. Thomy, P. Sass and H. Brötz-Oesterhelt designed the study. L. Reinhardt and D. Thomy conducted all experiments with the exception of intact-protein mass spectrometry. M. Lakemeyer and S. Sieber performed intact-protein mass spectrometry. J. Ortega contributed materials, plasmids and techniques. L. Reinhardt, D. Thomy, M. Lakemeyer, P. Sass and H. Brötz-Oesterhelt analysed data. P. Sass and H. Brötz-Oesterhelt supervised the study. L. Reinhardt, D. Thomy and P. Sass prepared figures and drafted the manuscript. All authors finalized and approved the manuscript. P. Sass and H. Brötz-Oesterhelt acquired funding.

## Funding

The authors appreciate funding by the *Deutsche Forschungsgemeinschaft* (DFG, German Research Foundation): Project-ID 174858087 (RTG 1708, project B5), Project-ID 398967434 (TRR 261, projects A01 and A02), as well as support by infrastructural funding from Cluster of Excellence EXC 2124 Controlling Microbes to Fight Infections, Project-ID 390838134). We further acknowledge financial support by the University of Duesseldorf/Research Centre Juelich (iGRASPseed fellowship to D. Thomy).

## Competing interests

The authors have declared that no competing interests exist.

## Notes

### Competing Interest Statement

The authors have declared no competing interest.

