## Supplementary Material for "Antibiotic acyldepsipeptides stimulate the *Streptomyces* Clp-ATPase/ClpP complex for accelerated proteolysis"

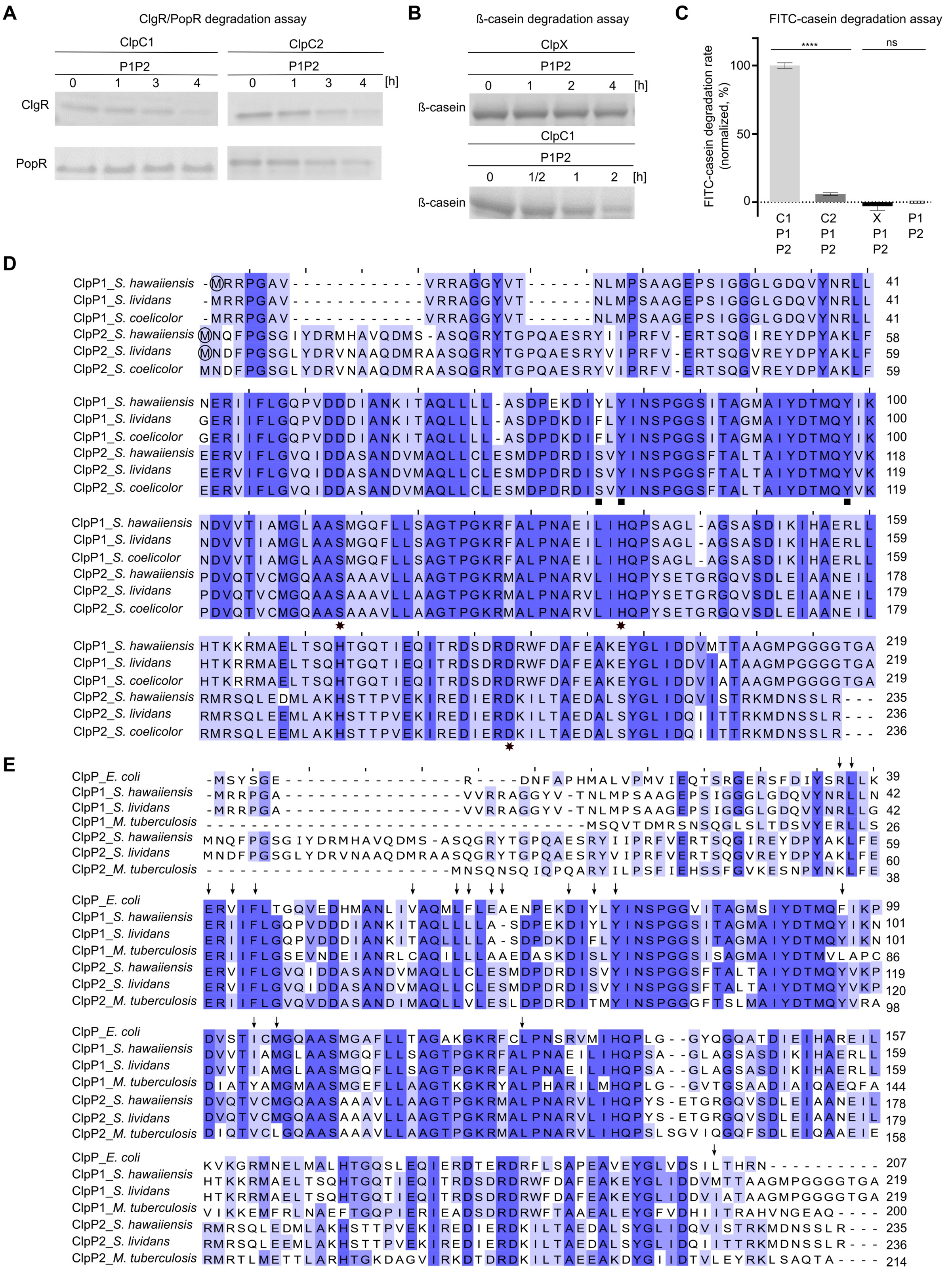


**Figure S1: Clp-ATPase-dependent protein degradation and amino acid sequence alignment of ClpP homologs from *Streptomyces spec., M. tuberculosis* and *E. coli.***

**A.** *In vitro* degradation of ClgR and PopR. As seen for ClpXP1P2 (Fig 1B), ClgR was also digested by ClpC1P1P2 and ClpC2P1P2, although with reduced efficiency. PopR was efficiently digested by ClpX (Fig 1B), still notably digested by ClpC2P1P2 but not by ClpC1P1P2. **B.** *In vitro* degradation of the model substrate ß-casein by ClpXP1P2 and ClpC1P1P2. ß-casein is not a substrate of the *Streptomyces* ClpXP1P2 protease, while it was degraded by ClpC1P1P2 and ClpC2P1P2 (Fig. 1B). All assays were performed in triplicates, and representative SDS-PAGE images are shown. **C.** FITC-casein degradation by ClpC1P1P2, ClpC2P1P2, ClpXP1P2 and ClpP1P2. Here, FITC-casein was significantly degraded by ClpC1P1P2, while ClpXP1P2, ClpC2P1P2 and ClpP1P2 showed no or only weak degradation of the substrate. Means of initial reaction kinetics are shown, normalized in %. Three biological replicates are shown each comprising three technical replicates. Statistics were performed with one-way ANOVA. P-values: ns > 0.05; **** ≤ 0.0001. Error bars indicate standard deviations. **D.** The multiple amino acid sequence alignment shows the positions of the catalytic triad residues (marked by stars) as well as of the three conserved aromatic residues within the hydrophobic pocket that were subject to site-directed mutagenesis in this study (marked by squares). Predicted GUG initiating codons (instead of AUG) are indicated with a circle. The alignment was generated with the online tool clustalΩ (https://www.ebi.ac.uk/Tools/msa/clustalo/) using ClpP protein sequences derived from *Streptomyces hawaiiensis* NRRL 15010, *Streptomyces lividans* TK24 and *Streptomyces coelicolor* A3(2). Sequence identities were computed using Jalview Software (74). In each column, the percentage of amino acid residues that agree with the consensus sequence are visualized from dark blue (> 80%) to marine (> 60%) to pale blue (> 40%). A comparison of the protein sequences of ClpP1 or ClpP2 from *S. hawaiiensis* and *S. lividans* using BLAST® (https://blast.ncbi.nlm.nih.gov/Blast.cgi) revealed sequence identities of 97% for ClpP1 and 95% for ClpP2 between the two species. **E.** Multiple sequence alignment of ClpP proteins from *Streptomyces spec.*, *M. tuberculosis* and *E. coli* illustrates amino acid residues, which are known to interact with the natural product ADEP1 (indicated by arrows) (34, 35). The alignment was generated with the online tool clustalΩ (https://www.ebi.ac.uk/Tools/msa/clustalo/). Sequence identities were computed using Jalview Software (74). In each column, the percentage of amino acid residues that agree with the consensus sequence are visualized from dark blue (> 80%) to marine (> 60%) to pale blue (> 40%).

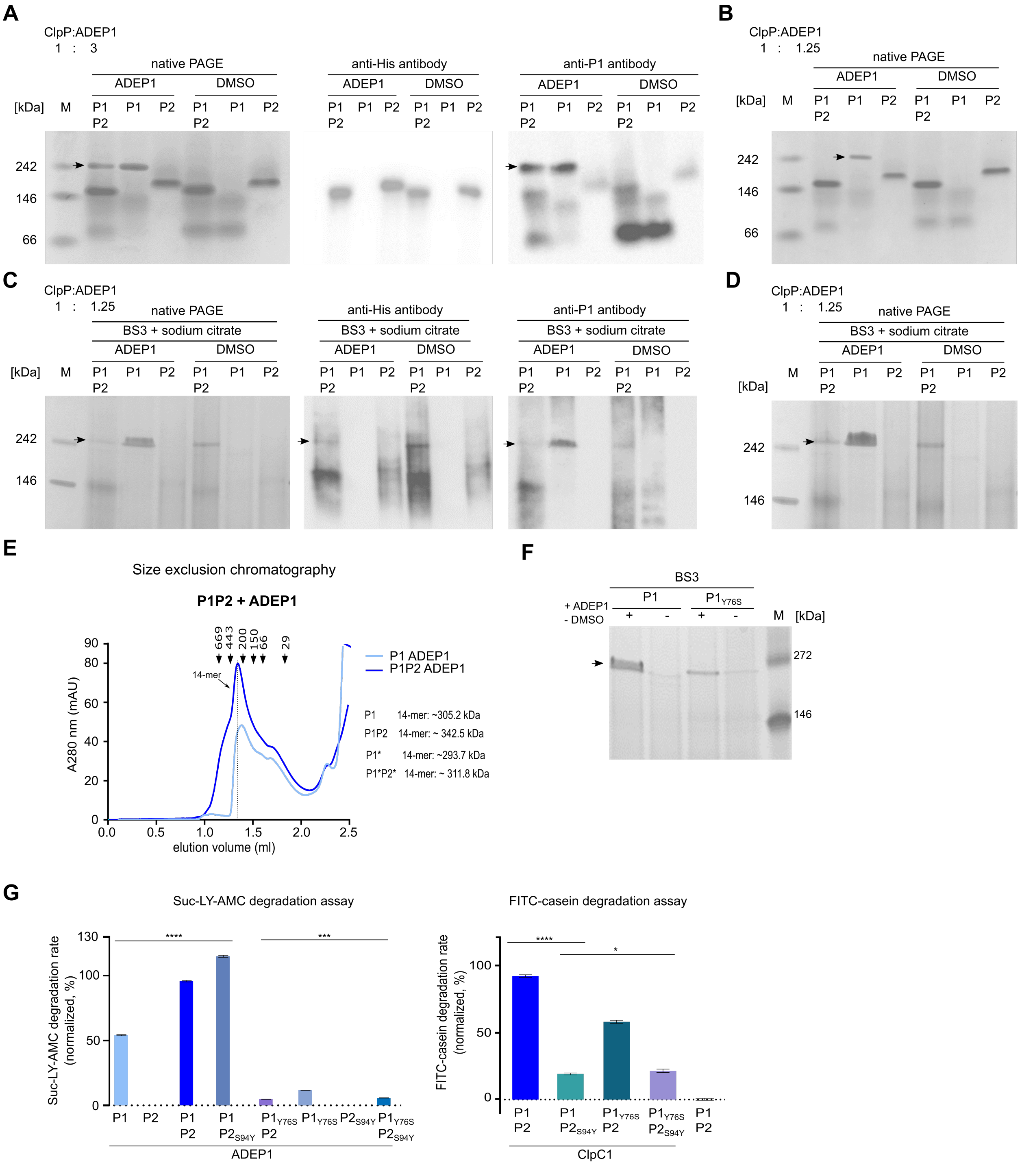


**Figure S2: Hetero-tetradecamer formation and peptidase/protease activities ClpP1 and ClpP2 wild-type proteins and ADEP/Clp-ATPase binding site mutants.**

**A.** Native PAGE, ClpP:ADEP ratio 1:3. On native PAGE gels, proteins bands with the size of tetradecamers (indicated by an arrow) were only observed in the presence of ADEP1 for samples comprising ClpP1P2 or ClpP1 alone. Immunoblotting using anti-ClpP1 or anti-6His antibodies identified those bands as ClpP1 homo-tetradecamers. Despite similar ClpP1 concentrations, the band of the ClpP1 homo-tetradecamer detected in the ADEP-ClpP1P2 sample appeared weaker on the native PAGE gel compared to the ADEP-ClpP1 sample. No protein bands with the size of tetradecamers were observed in the absence of ADEP1. However, immunoblotting using anti-6His antibodies detected heptameric ClpP2 in both samples ClpP1P2 and ClpP2 in the absence and presence ADEP1. **B.** Native PAGE, ClpP:ADEP ratio 1:1.25. Using decreased ADEP1 concentrations (ADEP1:ClpP1/ClpP2 ratio of 1.25:1), a tetradecamer band in the ClpP1P2 sample was no longer detectable, whereas ClpP1 homo-tetradecamers were still detected in the ClpP1 sample, although similar ClpP1 concentrations were used in both assays. **C**. Crosslinking experiments. In the absence and presence of ADEP1, the addition of the cross-linker BS3 and the complex stabilizing agent sodium citrate led to the detection of ClpP1P2 hetero-tetradecamers via native PAGE and subsequent immunoblotting. Remarkably, higher amounts of ClpP1 homo-tetradecamers were only detected in the ClpP1 sample in the presence of ADEP1 and absence of ClpP2. **D**. Increasing the amount of both ADEP and ClpP, while keeping their ratio constant, improved the detection of the tetradecameric protein band in the ClpP1P2 sample. **E.** Size exclusion chromatography of ClpP1P2 and ClpP1 alone and in the presence of ADEP1. Here, tetradecamers were detected in the ADEP-ClpP1P2 sample, which eluted slightly earlier than the tetradecameric fraction in the ADEP-ClpP1 sample. Hence, despite the rather small difference in size of approx. 18 kDa, the size-exclusion data allows for the differentiation of ADEP-bound ClpP1P2 hetero-tetradecamers versus and ADEP-bound ClpP1 homo-tetradecamers. The calculated mass of both ClpP1P2 and ClpP1 complexes, processed (indicated by an asterisk) or unprocessed are shown. The data is representative for three biological replicates. **F.** Using the cross-linker BS3, native PAGE showed decreased homo-tetradecamer formation of the ClpP1$_{Y76S}$ mutant compared to wild-type ClpP1 upon the addition of ADEP1. **G**. Suc-LY-AMC (left) and FITC-casein (right) degradation assays employing the mutant proteins ClpP1$_{Y76S}$ and ClpP2$_{S94Y}$. Our data indicates that Y76 of ClpP1 is important for the activation of ClpP1 by ADEP1, while S94 of ClpP2 is required for Clp-ATPase-ClpP2 interaction. In all assays, hydrolysis of Suc-LY-AMC and FITC-casein were recorded as an RFU increase over time. Mean values (normalized to %) of initial linear reaction kinetics are shown. Statistical analyses were performed with one-way ANOVA using three biological replicates each comprising three technical replicates. P-values: ns > 0.05; * ≤ 0.05; ** ≤ 0.01; ***≤ 0.001; ****≤ 0.0001. Error bars indicate standard deviations.

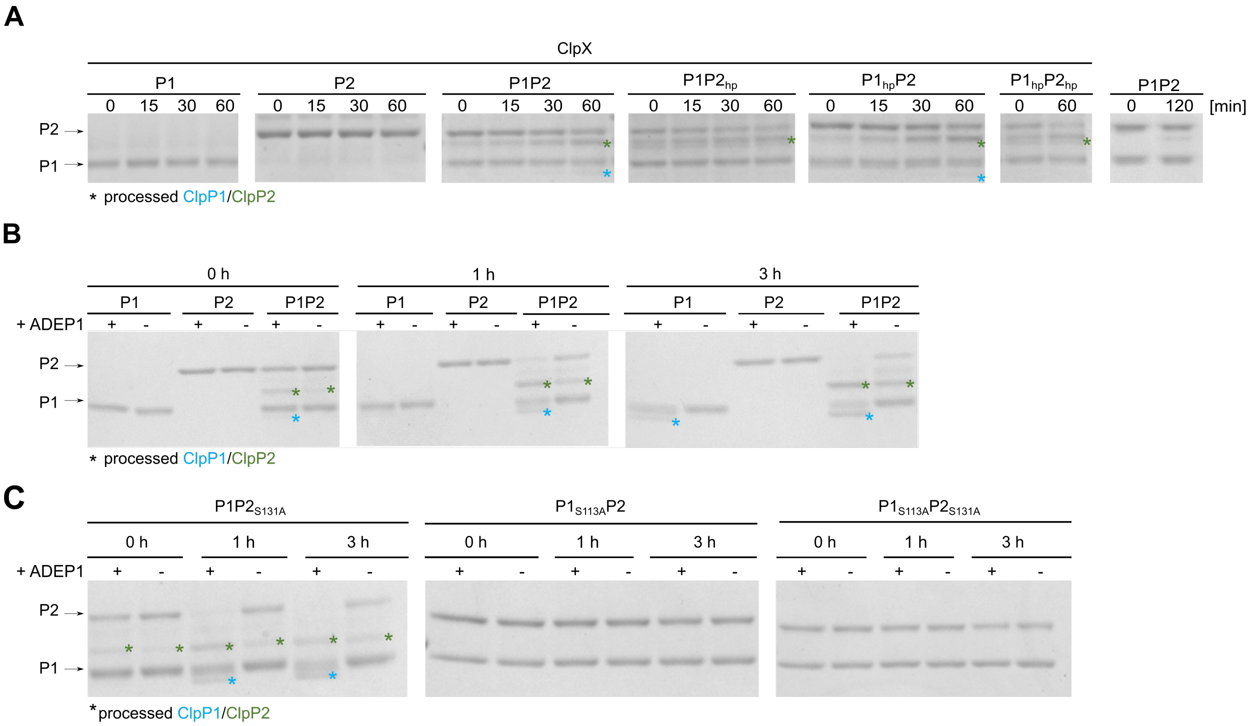


**D**

| | ClpP1 alone | ClpP2 alone | ClpP1P2 |
|---|---|---|---|
| Without activator | 0 | 0 | 0/+ |
| Clp-ATPase binding to ClpP2 | 0 | 0 | +/++ |
| ADEP1 binding to ClpP1 | + | 0 | ++/++ |

ClpP1; ClpP2 processing 0,+, ++

**Figure S3: ClpP1 and ClpP2 undergo processing reactions upon interaction with ADEP or Clp-ATPases.**

**A.** ClpP1 and ClpP2 are processed when used in combination and in the presence of ClpX. Processed proteins are marked by coloured asterisks (ClpP1*, blue; ClpP2*, green). Processing of ClpP1 depends on activation by Clp-ATPase binding to ClpP2 and is prevented in the hydrophobic patch mutant ClpP2$_{hp}$. In contrast, ClpP2 processing solely depends on hetero-tetradecamer formation, but not on binding and activation by a Clp-ATPase. **B.** ADEP1 induces processing of ClpP1 and accelerates processing of ClpP2. Samples were incubated in the absence or presence of ADEP1 over time. Processing of ClpP1 requires the presence of an activator (here ADEP1). The processing of ClpP1 occurred in samples with mixed ClpP1P2 and more slowly by ClpP1 alone as indicated by blue asterisks. Thus, ADEP, in contrast to a Clp-ATPase, can stimulate ClpP1 processing in the absence of ClpP2. Unlike ClpP1, processing of ClpP2 solely depends on hetero-tetradecamer formation, but not on binding and activation by ADEP1. However, processing of ClpP2 is notably accelerated in the presence of ADEP1. **C.** Processing of ClpP1 and ClpP2 depends on the catalytic function of ClpP1. Processing of both ClpP1 and ClpP2 was prevented in samples containing the catalytic triad mutant ClpP1$_{S113A}$, whereas ClpP2$_{S131A}$ had no effect on the processing reactions. All assays were performed at least in triplicates. DMSO was used in control reactions. Processed forms of ClpP1 and ClpP2 are indicated by coloured asterisks (ClpP1*, blue; ClpP2*, green). Representative SDS-PAGE images are shown. **D.** Overview of processing reactions.

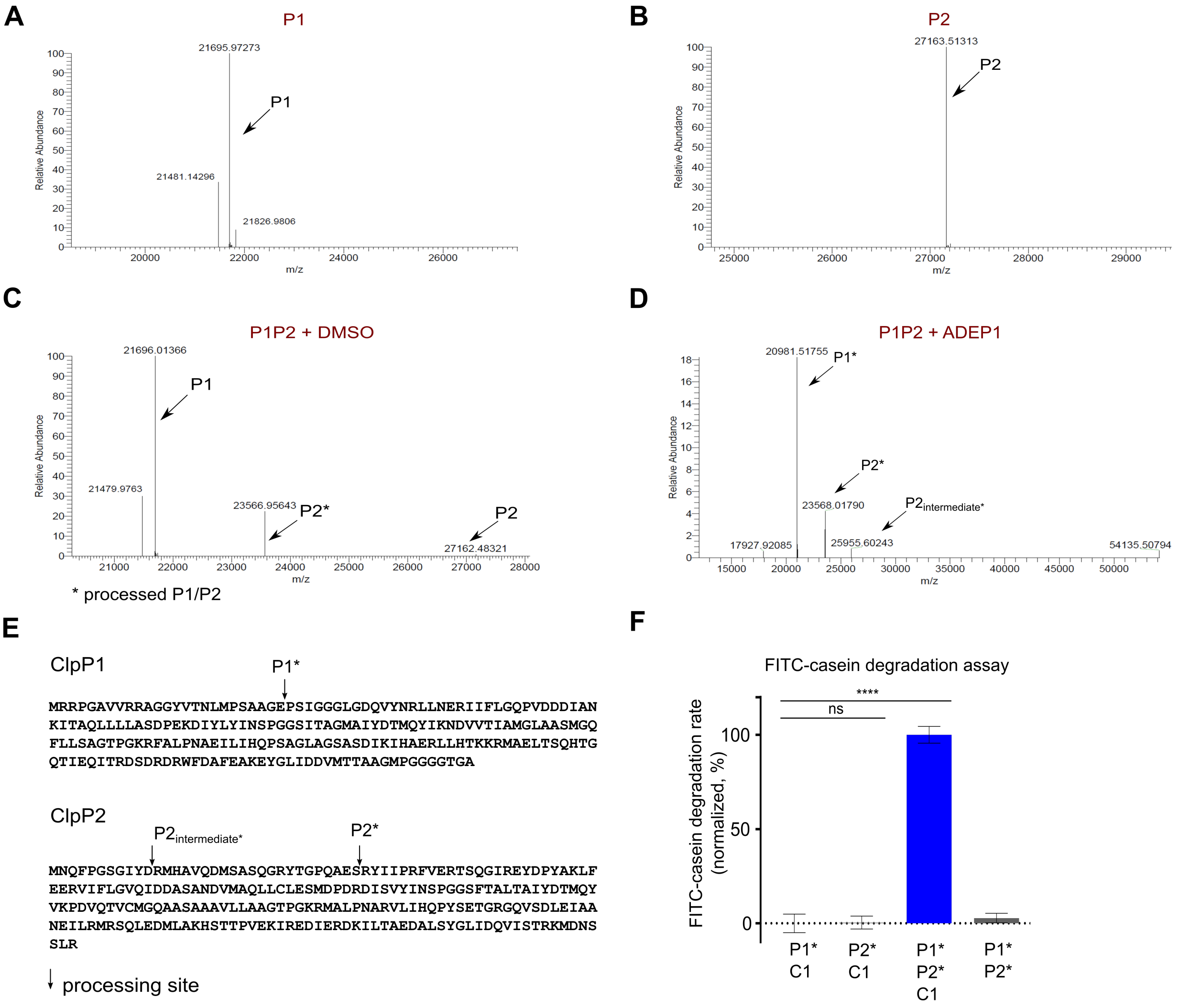


**Figure S4: Intact protein mass spectrometry and proteolytic activity of processed variants of ClpP1 and ClpP2.** **A-D.** Intact protein mass spectrometry of (A) ClpP1 alone, (B) ClpP2 alone, (C) ClpP1 + ClpP2 + DMSO, and (D) ClpP1 + ClpP2 + ADEP1 verified that ClpP2 is processed by ClpP1 in the absence and presence of ADEP1 (indicated by P2*), whereas ClpP1 processes itself only in the presence of ADEP1 (indicated by P1*). Our data show that ClpP2 is processed with one intermediate processing step ($P2_{intermediate}$*), finally resulting in mature ClpP2 (P2*). **E.** Amino acid sequences of ClpP1 and ClpP2 indicating the processing sites according to intact protein mass spectrometry. ClpP2 is first processed between amino acids D11 and R12, resulting in an intermediately processed ClpP2 ($P2_{intermediate}$*) that starts with the amino acids RMHAVQ. The second processing site of ClpP2 is located between amino acids S33 and R34, resulting in mature ClpP2 (P2*) starting with the amino acid sequence RYIIPR. The detected mass for processed ClpP1 (P1*) indicates the processing site to be located between the amino acids G24 and P25. P1* was only detected in the presence of ADEP1. **F.** To investigate the effect of processing on the proteolytic activity, we cloned and purified the processed variants ClpP1* and ClpP2* and used them in FITC-casein degradation assays in the absence or presence of ClpC1. As observed before, when unprocessed ClpP1 and ClpP2 were subjected to the same assay, proteolytic digestion of FITC-casein did only occur in the presence of both ClpP isoforms, (here ClpP1* and ClpP2*) plus ClpC1, proving that the processed forms still rely on hetero-tetradecamer formation and the presence of a Clp-ATPase for proteolytic activity (for processing induced by ADEP compare Fig S3). Hydrolysis of FITC-casein was monitored by an increase in the fluorescence signal (RFUs) over time. Mean values (normalized in %) of the initial linear reaction kinetics are given. P-values were calculated with one-way ANOVA using three biological replicates each comprising three technical replicates. P-values: ns > 0.05; **** ≤ 0.0001. Error bars indicate standard deviations.

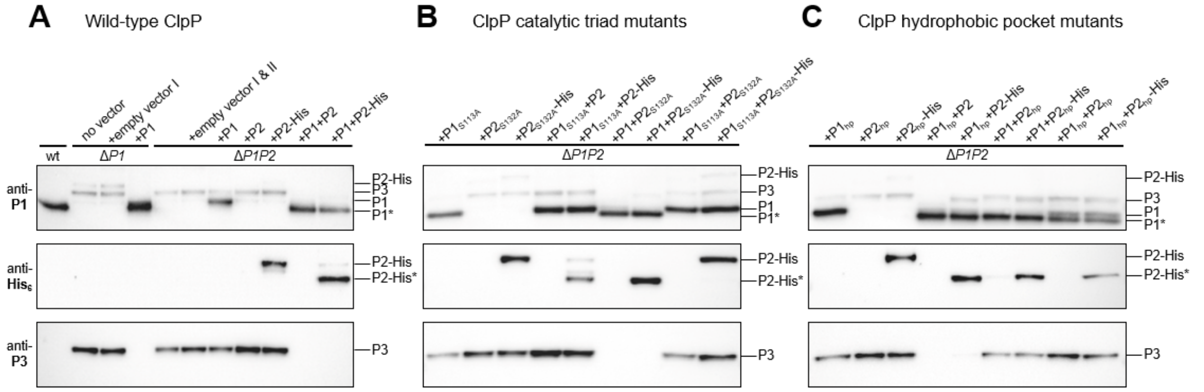


**Figure S5: Whole cell studies of Clp protease activity using wild-type *S. lividans* and *clpP1*/*clpP2* mutants.**

Immunoblotting of cell lysates from wild-type *S. lividans* cells and *clpP1*/*clpP2* deletion mutants (**A**), catalytic triad mutants (**B**) or hydrophobic pocket mutants (**C**), as well as respective complemented strains, using anti-ClpP1, anti-ClpP3 and anti-His$_6$ antibodies. Both SlClpP1 and SlClpP2 are required to build a functional, heteromeric Clp protease system in whole cells, which strictly depends on a functional catalytic triad of SlClpP1 for proteolytic activity and on a functional hydrophobic pocket of SlClpP2 for the interaction with corresponding Clp-ATPases (both indicated by the loss of PopR degradation and concomitant expression of SlClpP3 in mutants harbouring either ClpP1$_{S113A}$ or ClpP2$_{hp}$, respectively). All assays were performed at least in triplicates. Representative Western Blot images are shown. Noteworthy, in the cell extract of the strain Δ*SlP1P2* expressing the proteins SlClpP1$_{S113A}$ and SlClpP2-His, slight signals of processed forms of SlClpP2-His were detectable, which did not appear in the absence of SlClpP1$_{S113A}$ and were also not visible in our *in vitro* experiments. Thus, in the more complex and optimized whole cell environment the interaction with SlClpP1$_{S113A}$ might permit a weak catalytic activity of SlClpP2, sufficient for self-processing under appropriate conditions. However, in none of our assays, neither *in vivo* nor *in vitro,* we could detect substrate degradation activity of the ClpP2 active sites within the *Streptomyces* ClpP1P2 complex. Interestingly, the hydrophobic pocket mutations did not affect the processing of ClpP1 and ClpP2, indicating that processing of ClpP1 does not strictly rely on Clp-ATPase binding to ClpP2 in the living cell, representing a situation with additional, potentially necessary interacting factors present, in contrast to the *in vitro* situation that is limited to a defined set of proteins.

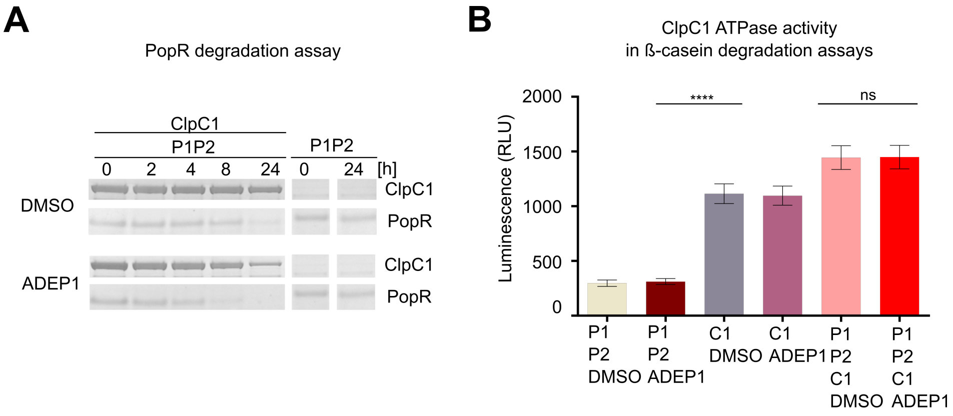


**Figure S6: ClpC1 protease activity and ATP consumption in the presence of ADEP1.**

**A.** ClpC1 is self-digested by ClpC1P1P2 after prolonged incubation. *In vitro* protein degradation assays employing purified Clp proteins as indicated and PopR as a substrate. The data show that the digestion of PopR by ClpC1P1P2 is accelerated by ADEP1 compared to the DMSO control. Of note, in the presence of ADEP1 and after PopR has been fully digested, ClpC1 is also slowly degraded in ClpC1P1P2 reactions. All assays were performed in triplicates and representative SDS-PAGE images are shown. DMSO was used in control reactions. **B.** ClpC1-mediated ATPase activity in casein degradation assays. To answer the question, whether the observed increase of Clp-ATPase/ClpP1P2 activity in the presence of ADEP1 was due to a more rapid utilization of ATP by the AAA+ ATPases, we directly measured the turn-over of ATP to ADP by ClpC1 in casein degradation assays. Here, ATP turn-over was not increased in the presence of ADEP (compared to the DMSO control), neither for ClpCP1P2 nor ClpC1 alone. Hence, stimulated ATPase/ClpP1P2 activity in the presence of ADEP1 is most probably unrelated to ATP turn-over. Instead, ADEP1 binding may affect the overall structural network of the ClpP1P2 complex, including stabilization in the extended conformation and the adjustment of the active site residues as well as pore opening, as was previously described for ADEP-activated ClpP in other bacteria (34, 35, 38). Mean values of three biological replicates are shown. Statistics were performed with one-way ANOVA. P-values: ns > 0.05; **** ≤ 0.0001. Error bars indicate standard deviations.

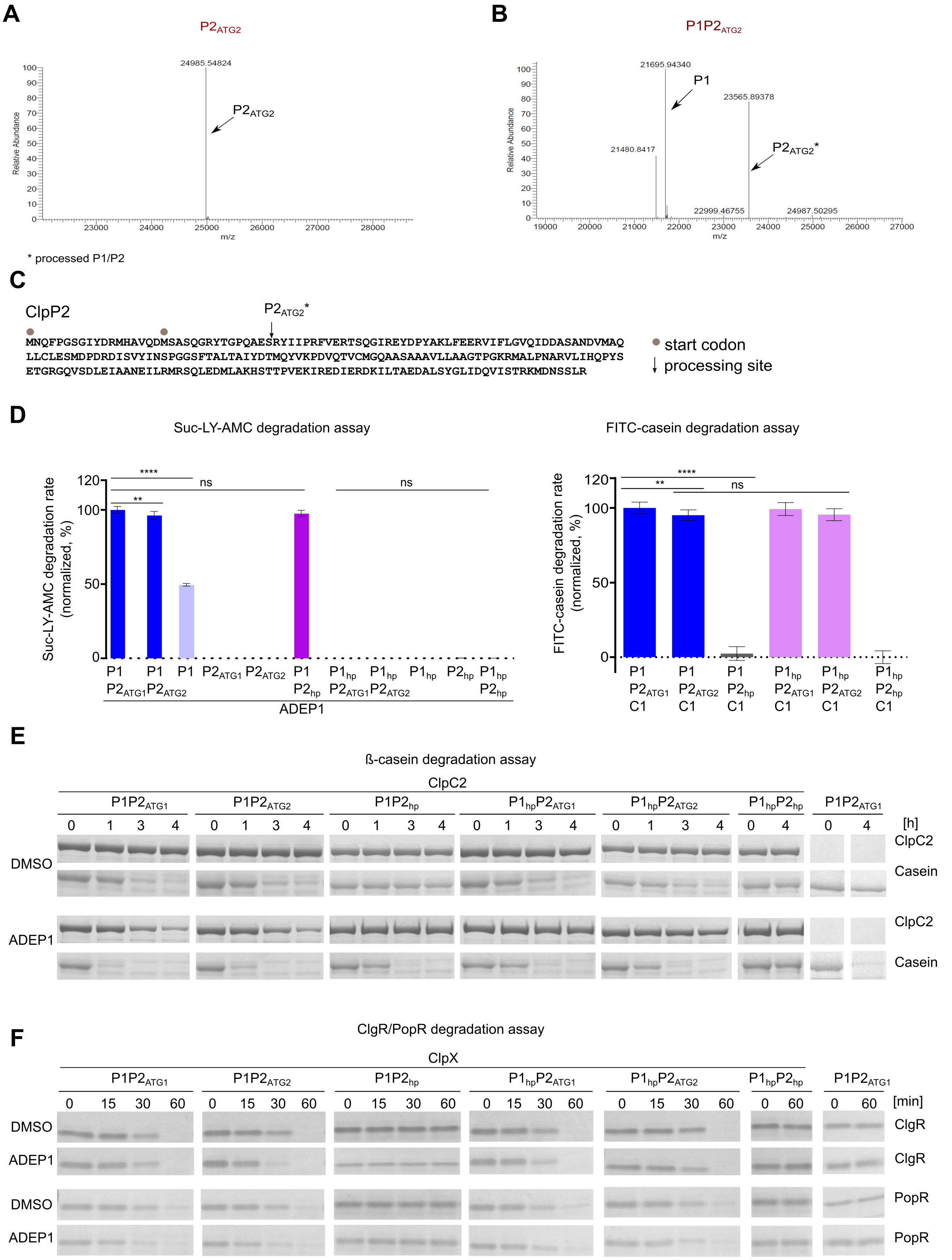


**Figure S7: Intact protein mass spectrometry and processing analysis of Clp2$_{ATG2}$, as well as peptidase/protease activities of Clp2$_{ATG1}$ versus Clp2$_{ATG2}$ and hydrophobic pocket mutants.**

**A-B.** Intact protein mass spectrometry of (A) ClpP2$_{ATG2}$ and (B) ClpP1 + ClpP2$_{ATG2}$. In principle, ClpP2 may also be expressed from a putative second start codon (here indicated by ClpP2$_{ATG2}$). We therefore cloned and expressed ClpP2$_{ATG2}$ using the putative second start codon and tested its processing behaviour using intact protein mass spectrometry compared to ClpP2. Our data show that the putative second start codon of ClpP2 results in the same processed mature protein compared to expression from the first start codon, as it is shown in Fig S4. **C.** Amino acid sequence of full-length ClpP2 highlighting the first and putative second start codon (grey dots) as well as the resulting processing site deduced from intact protein mass spectrometry. The detected mass for processed ClpP2$_{ATG2}$* indicates the same processing site between the amino acids S33 and R34, resulting in a protein that starts with RYIIPR, similar to processed ClpP2*. **D.** Suc-LY-AMC (left) and FITC-casein (right) degradation assays showing peptidase and protease activity, respectively, of ClpP2$_{ATG1}$ compared to ClpP2$_{ATG2}$. Clp wild-type and mutant proteins as well as ADEP1 were used as indicated. The data corroborates our results shown above regarding the role of ClpP1 in ADEP-mediated proteolysis as well as of ClpP2 for Clp-ATPase binding. In all assays, hydrolysis of Suc-LY-AMC and FITC-casein were recorded as an RFU increase over time. Mean values (normalized to %) of initial linear reaction kinetics are shown. Statistical analyses were performed with one-way ANOVA using three biological replicates each comprising three technical replicates, with the exception that two biological replicates were used in Suc-LY-AMC degradation assays employing the hydrophobic pocket mutants ClpP1$_{hp}$ and ClpP2$_{hp}$. P-values: ns > 0.05; ** ≤ 0.01; **** ≤ 0.0001. Error bars indicate standard deviations. **E-F.** *In vitro* ß-casein, ClgR, or PopR degradation assays employing purified Clp proteins and ADEP1/DMSO as indicated. Corroborating our results above, the use of either ClpP2$_{ATG1}$ or ClpP2$_{ATG2}$ resulted in comparable activities in these assays. Further noteworthy, accelerated substrate degradation in the presence of ADEP1 also occurred in samples using ClpP2$_{ATG2}$. All assays were performed at least in triplicates and representative SDS-PAGE images are shown. DMSO was used in control reactions.

**Table S1. Bacterial strains and Plasmids.**

| Strain /plasmid | Relevant characteristic(s)/genotype | Ref. /Source |
|---|---|---|
| *E. coli* | | |
| K-12 JM109 | subcloning host | (75) |
| DH5α | subcloning host | Thermo Fisher Scientific |
| SG1146a | BL21(DE3) *ClpP::cam* | (72) |
| ET12567 pUB307 | *F-dam-13::Tn9 dcm-6 hsdM hsdR zjj-202::Tn10 recF143 galK2 galT22 ara-14 lacY1 xyl-5 leuB6 thi-1 tonA31 rpsL136 hisG4 tsx-78 mtl-1 glnV44; pUB307;* $Cm^R$, $Kan^R$ | (76) (77) |
| *S. lividans* | | |
| TK24 | *str-6;* $SLP2^-$, $SLP3^-$ | (78) |
| Δ*clpP1* | *str-6;* $SLP2^-$, $SLP3^-$; Δ*clpP1* | this study |
| Δ*clpP1clpP2* | *str-6;* $SLP2^-$, $SLP3^-$; Δ*clpP1clpP2* | this study |
| Plasmids | | |
| pET11a | vector for the expression of native protein expression | Novagen |
| pET22b | vector for the expression of C-terminal His6-fusion protein | Novagen |
| pETDUET-1 | vector for the co-expression of two target genes | Novagen |
| pET28aShclpP1 | pET28a + ORF CEB94_14110 (*S. hawaiiensis clpP1*) | this study |
| pET21bShclpP2 | pET21b + ORF CEB94_14105 (*S. hawaiiensis clpP2*) | this study |
| pET11aShclpP1$_{ATG2}$ | pET11a + ORF CEB94_14110 (*S. hawaiiensis clpP1*) | this study |
| pET11aShclpP2 | pET11a + ORF CEB94_14105 (*S. hawaiiensis clpP2*) | this study |
| pET22bShclpP1$_{ATG2}$-His6 | pET22b + ORF CEB94_14110 (*S. hawaiiensis clpP1*) | this study |
| pET22b**Nco*I-ShclpP2-His6 | pET22b**Nco*I + ORF CEB94_14105 (*S. hawaiiensis clpP2*) | this study |
| pET22b**Nco*I-ShclpP2$_{ATG2}$-His6 | pET22b**Nco*I + ORF CEB94_14105 (*S. hawaiiensis clpP2*) | this study |
| pET22bShclpP1*-His6 | pET22b + ORF CEB94_14110 (*S. hawaiiensis clpP1*) | this study |
| pET22b**Nco*I-ShclpP2*-His6 | pET22b + ORF CEB94_14105 (*S. hawaiiensis clpP2*) | this study |
| pET11aShclgR-N-His6 | pET11a + ORF CEB94_30145 (*S. hawaiiensis clgR*) | this study |
| pET11aShpopR-N-His-6 | pET11a + MT943519 (*S. hawaiiensis popR*) | this study |
| pETDUETShclpP1$_{ATG2}$clpP2-His6 | pETDUET-1 + ORF CEB94_14110 + 14105 (*S. hawaiiensis clpP1+P2*) | this study |
| pET22b**Nco*I-ShclpX-His6 | pET22b**Nco*I + ORF CEB94_14100 (*S. hawaiiensis clpX*) | this study |
| pET22b**Nco*I-ShclpC1-His6 | pET22b*NcoI + ORF CEB94_23085 (*S. hawaiiensis clpC1*) | this study |
| pET22bShclpC2-His6 | pET22b + ORF CEB94_33910 (*S. hawaiiensis clpC2*) | this study |
| pET22bShclpP1$_{S113A}$ | pET22bshclpP1$_{ATG2}$-His6 carrying aa mutation S113A in the *S. hawaiiensis clpP1* gene | this study |

| | | |
|---|---|---|
| pET22bShclpP2$_{S131A}$ | pET22b**Nco*I-ShclpP2-His6 carrying aa mutation S131A in the *S. hawaiiensis clpP2* gene | this study |
| pET11aShclpP1$_{hp}$ | pET11aShclpP1$_{ATG2}$ carrying aa mutation Y76V, Y78V, Y98V in the *S. hawaiiensis clpP1* gene | this study |
| pET22bShclpP2$_{hp}$ | pET22b**Nco*I-ShclpP2$_{ATG2}$-His6 carrying aa mutation S94A, Y96V, Y116V in the *S. hawaiiensis clpP2* gene | this study |
| pET11aShclpP1$_{Y76SATG2}$ | pET11aShclpP1$_{ATG2}$ carrying aa mutation Y76S in the *S. hawaiiensis clpP1* gene | this study |
| pET22b**Nco*I-ShclpP2$_{S94YATG2}$-His6 | pET22b**Nco*I-ShclpP2$_{ATG2}$-His6 carrying aa mutation S94Y in the *S. hawaiiensis clpP2* gene | this study |
| pGM-GUS | temperature-sensitive *Streptomyces* shuttle vector *aac(3)IV, oriT, $P_{ermE}$_gusA, $rep_{ts}$* | Günther Muth, Tübingen |
| pGM-GUS-Xba | based on pGM-GUS, introduction of an XbaI restriction site by site-directed mutagenesis | this study |
| pGM-GUS-clpP1 | knockout vector for *S. lividans clpP1* | this study |
| pGM-GUS-clpP1clpP2 | knockout vector for *S. lividans clpP2clpP2* | this study |
| pIJ12551 | ΦC31-integrative Streptomyces shuttle vector, protein expression under *ermE** promotor | (79) |
| pIJ12551clpP1 | constitutive protein expression of SlClpP1 | this study |
| pIJ12551clpP1$_{S113A}$ | constitutive protein expression of SlClpP1 with the following mutation(s): S113A | this study |
| pIJ12551clpP1$_{hp}$ | constitutive protein expression of SlClpP1 with the following mutation(s): Y76V, Y78V, Y98V | this study |
| pIJ12551clpP1clpP2 | constitutive protein expression of SlClpP1ClpP2 | this study |
| pIJ10257 | ΦBT1-integrative Streptomyces shuttle vector, protein expression under *ermE** promotor | (80) |
| pIJ10257clpP2 | constitutive expression of SlClpP2 | this study |
| pIJ10257clpP2-His | constitutive expression of SlClpP2 with an N-terminal 6xHis-tag | this study |
| pIJ10257clpP2$_{S132A}$ | constitutive protein expression of SlClpP2 with the following mutation(s): S132A | this study |
| pIJ10257clpP2-His$_{S132A}$ | constitutive protein expression of SlClpP2 with an N-terminal 6xHis-tag with the following mutation(s): S132A | this study |
| pIJ10257clpP2$_{hp}$ | constitutive protein expression of SlClpP2 with the following mutation(s): S95A, Y97V, Y117V | this study |
| pIJ10257clpP2-His$_{hp}$ | constitutive protein expression of SlClpP2 with an N-terminal 6xHis-tag with the following mutation(s): S95A, Y97V, Y117V | this study |

**Table S2. Primer used in this study.** Restriction sites are underlined.

| Plasmid | | Forward (F)/Reverse (R) oligo (5'- 3' direction) | Template |
|---|---|---|---|
| pET11aShclpP1$_{ATG2}$ | | F: aaacatatgacgaatctgatgccctcagc | pET28aShclpP1 |
| | | R: aaaggatcctcaggccccggtgccgc | |
| pET11aShclpP2 | | F: aaacatatgaaccagttccccggcag | *S. hawaiiensis* genomic DNA |
| | | R: aaaggatcctcagcgcaggctcgagttgtc | |
| pET22bShclpP1$_{ATG2}$-His6 | | F: aaacatatgacgaatctgatgccctcagc | pET28aShclpP1 |
| | | R: taaactcgagggccccggtgccgcc | |
| pET22b**Nco*I-ShclpP2-His6 | | F: aaacatatgaaccagttccccggcag | *S. hawaiiensis* genomic DNA |
| | | R: aaaccatgggcgcaggctcgagttgtc | |
| pET22b**Nco*I-ShclpP2$_{ATG2}$-His6 | | F: aaacatatgagcgcctcccagggc | pET21bShclpP2 |
| | | R: aaaccatgggcgcaggctcgagttgtc | |
| pET22bShclpP1*-His6 | | F: aaacatatgccttccatcggtggtggcctcggt | pET22bShclpP1$_{ATG2}$-His6 |
| | | R: taaactcgagggccccggtgccgcc | |
| pET22b**Nco*I-ShclpP2*-His6 | | F: aaacatatgcgctacatcattccccgcttc | pET22b**Nco*I-ShclpP2-His6 |
| | | R: aaaccatgggcgcaggctcgagttgtc | |
| pET11aShclgR-N-His6 | | F: aaacatatgcaccaccaccaccaccacattctgctccgtcgcctgctgggtgacgtg | *S. hawaiiensis* genomic DNA |
| | | R: taaggatcctcacgcggcgacgacgtccactgc | |
| pET11aShpopR-N-His-6 | | F: aaacatatgcaccaccac caccaccacaccagccacctgccgaacgaagcccgagtc | *S. hawaiiensis* genomic DNA |
| | | R: taaggatcctcaggcggccaggcacattccgtcgtg | |
| pETDUETShclpP1$_{ATG2}$clpP2-His6 | clpP1 | F: aaacatatgacgaatctgatgccctcagc | pET11aShclpP1$_{ATG2}$ |
| | | R: aaaggtacctcaggccccggtgcc | |
| | clpP2 | F: aaaccatgggaaaccagttccccggcagcg | pET22b**Nco*I-ShclpP2-His6 |
| | | R: aaaaagctttcagtggtggtggtggtggtggcgcaggctcgagttgtccatcttcc | |
| pET22b**Nco*I-ShclpX-His6 | | F: aaacatatggcacgcatcggtgacggcg | genomic DNA *S. hawaiiensis* |
| | | R: aaaccatggggccgtcttctgctcccccgg | |
| pET22b**Nco*I-ShclpC1-His6 | | F: aagcatatgttcgagaggttcaccgacc | genomic DNA *S. hawaiiensis* |
| | | R: aaaccatggggcgtccttgctcaggttcggg | |
| pET22bShclpC2-His6 | | F: aaacatatgagcagcggcttcaccagc | *S. hawaiiensis* genomic DNA |
| | | R: t aaactcgagtcggggcacggtactgaacg | |
| pET22bShclpP1$_{S113A}$ | | F: gggcctggcagccgcgatgggccagttc | pET22bShclpP1$_{ATG2}$-His6 |
| | | R: gaactggcccatcgcggctgccaggccc | |
| pET22bShclpP2$_{S131A}$ | | F: ccaggcggccgccgccgccgccgtc | pET22b**Nco*I-ShclpP2-His6 |
| | | R: gacggcggcggcggcggccgcctgg | |
| pET11aShclpP1$_{hp}$ | | F: ggagaaggacatcgtcctggtcatcaacagccccggc | pET11aShclpP1$_{ATG2}$ |
| | | R: cctcttcctgtagcaggaccagtagttgtcggggccg | |

| | | |
|---|---|---|
| | F: gacaccatgcaggtcatcaagaacgac | |
| | R: gtcgttcttgatgacctgcatggtgtc | |
| pET22bShclpP2$_{hp}$ | F: cgaccgtgacatcgcggtggtcatcaacagccccggc | pET22b**Nco*I-ShclpP2$_{ATG2}$-His6 |
| | R: gctggcactgtagcgccaccagtagttgtcggggccg | |
| | F: gacacgatgcaggtcgtgaagccggac | |
| | R: gtccggcttcacgacctgcatcgtgtc | |
| pET11aShclpP1$_{Y76S}$ | F: gagaaggacatctccctgtacatcaacag | pET11aShclpP1$_{ATG2}$ |
| | R: ctgttgatgtacagggagatgtccttctc | |
| pET22bShclpP2$_{S94YATG2}$-His6 | F: gaccgtgacatctacgtgtacatcaac | pET22b**Nco*I-ShclpP2$_{ATG2}$-His6 |
| | R: gttgatgtacacgtagatgtcacggtc | |
| pGM-GUS-Xba | F: cccgcgcccagtccgagctctagacggcgcttttcacctggc | pGM-GUS |
| | R: gccaggtgaaaagcgccgtctagagctcggactgggcgcggg | |
| pGM-GUS-clpP1 | F1: ctgcagacgcgtcgacgtcatatgacatcacggagctgaag | pGM-GUS-Xba, |
| | R1: cgatcgtctgcacgtatccacctgctcg | *S. lividans* TK24 genomic DNA |
| | F2: tggatacgtgcagacgatcgagcagatc | |
| | R2: ccacggcgatatcggatccatatgccagcaggaggatgttgg | |
| pGM-GUS-clpP1clpP2 | F1: tggctccaattgtacatcggatccatatggacatcacggagctgaag | pGM-GUS-Xba, |
| | R1: tggtgatgatctggtccacgtatccacctgctcg | *S. lividans* TK24 genomic DNA |
| | F2: aggtggatacgtggaccagatcatcaccaccc | |
| | R2: agcttctgcagacgcgtcgacgtcatatggaatcccgggggatcagc | |
| pIJ12551clpP1 | F: ggaattccatatgacgaatctgatgccctcag | pIJ12551, |
| | R: atagtttagcggccgctcaggcgcccgtgccgccgc | *S. lividans* TK24 genomic DNA |
| pIJ12551clpP1$_{S113A}$ | F: gcgatgggtctcgcggccGccatgggacagttcctgc | pIJ12551clpP1 |
| | R: gcaggaactgtcccatggcggccgcgagacccatcgc | |
| pIJ12551clpP1$_{hp}$ | F1: gacccggacaaggacatcGTcctgGTcatcaacagcccgggcgg | pIJ12551clpP1 |
| | R1: ccgcccgggctgttgatgaccaggacgatgtccttgtccgggtc | |
| | F2: gatctacgacaccatgcagGTcatcaagaacgacgtggtg | |
| | R2: caccacgtcgttcttgatgacctgcatggtgtcgtagatc | |
| pIJ12551clpP1clpP2 | F: ggaattccatatgacgaatctgatgccctcag | pIJ12551, |
| | R: atagtttagcggccgcctagcggagagaggagttgtc | *S. lividans* TK24 genomic DNA |
| pIJ10257clpP2 | F: ggaattccatatgaacgacttccccggcagcg | pIJ10257, |
| | R: cccaagcttctagcggagagaggagttgtc | *S. lividans* TK24 genomic DNA |
| pIJ10257clpP2-His | F: ggaattccatatgaacgacttccccggcagcg | pIJ10257, |
| | R: cccaagcttctaatgatgatgatgatgatggcggagagaggagttgtc | *S. lividans* TK24 genomic DNA |
| pIJ10257clpP2$_{S131A}$ | F: gtctgcatgggccaggccgccgccgccgccgccgtcctgctgg | pIJ10257clpP2 |
| | R: ccagcaggacggcggcggcggcggcggcctggcccatgcagac | |

| | | |
|---|---|---|
| pIJ10257clpP2-His$_{S132A}$ | F: gtctgcatgggccaggccgccgccgccgccgccgtcctgctgg | pIJ10257clpP2-His |
| | R: ccagcaggacggcggcggcggcggcggcctggcccatgcagac | |
| pIJ10257clpP2$_{hp}$ | F1: cccgaccgggacatcgcggtcgtcatcaacagccccgg | pIJ12057clpP2 |
| | R1: ccggggctgttgatgacgaccgcgatgtcccggtcggg | |
| | F2: ctacgacacgatgcaggtcgtgaagccggacgtccagac | |
| | R2: gtctggacgtccggcttcacgacctgcatcgtgtcgtag | |
| pIJ10257clpP2-His$_{hp}$ | F1: cccgaccgggacatcgcggtcgtcatcaacagccccgg | pIJ12057clpP2-His |
| | R1: ccggggctgttgatgacgaccgcgatgtcccggtcggg | |
| | F2: ctacgacacgatgcaggtcgtgaagccggacgtccagac | |
| | R2: gtctggacgtccggcttcacgacctgcatcgtgtcgtag | |
